# TAK1-mediated phosphorylation of PLCE1 represses PIP2 hydrolysis to impede esophageal squamous cancer metastasis

**DOI:** 10.1101/2024.03.22.586256

**Authors:** Qianqian Ju, Wenjing Sheng, Meichen Zhang, Jing Chen, Liucheng Wu, Xiaoyu Liu, Wentao Fang, Hui Shi, Cheng Sun

## Abstract

TAK1 is a serine/threonine protein kinase that is a key regulator in a wide variety of cellular processes. However, the functions and mechanisms involved in cancer metastasis are still not well understood. Here, we found that TAK1 knockdown promoted esophageal squamous cancer cell (ESCC) migration and invasion, whereas TAK1 overexpression resulted in the opposite outcome. These *in vitro* findings were recapitulated *in vivo* in a xenograft metastatic mouse model. Mechanistically, co-immunoprecipitation and mass spectrometry demonstrated that TAK1 interacted with phospholipase C epsilon 1 (PLCE1), and phosphorylated PLCE1 at serine 1060 (S1060). Functional studies revealed that phosphorylation at S1060 in PLCE1 resulted in decreased enzyme activity, leading to the repression of PIP2 hydrolysis. As a result, the degradation products of PIP2 including diacylglycerol (DAG) and inositol IP3 were reduced, which thereby suppressed signal transduction in the axis of PKC/GSK-3β/β-Catenin. Consequently, expression of cancer metastasis-related genes was impeded by TAK1. Overall, our data indicate that TAK1 plays a negative role in ESCC metastasis, which depends on the TAK1 induced phosphorylation of PLCE1 at S1060.

## Introduction

Esophageal cancer (EC) is the seventh most common malignancy worldwide, with more than 600,000 new cases diagnosed annually (Sung, Ferlay et al., 2021). The primary histological subtypes of EC include esophageal squamous cell carcinoma (ESCC) and esophageal adenocarcinoma (He, Xu et al., 2021, Lagergren, Smyth et al., 2017). ESCC is a major subtype of EC in China, accounting for more than 90% of EC (He et al., 2021). The early diagnosis rate of ESCC is very low owing to the lack of specific biomarkers. Thus, most patients are found to be in locally advanced or metastatic stages when they are diagnosed. The 5-year survival rate is approximately 15-25% (Pennathur, Gibson et al., 2013). In recent decades, although therapeutic strategies have made great progress towards ESCC, recurrence and metastasis still remain significant challenges (Mao, Zeng et al., 2021). Therefore, there is an urgent need to deeply understand the mechanisms involved in cell metastasis, and thus try to explore new druggable targets for treating ESCC.

Transforming growth factor β-activated kinase 1 (TAK1), a member of the mitogen-activated protein kinase (MAPK) kinase kinase (MAP3K) family, is encoded by the gene *Map3k7*. As a serine/threonine protein kinase, TAK1 has been shown to play an integral role in inflammatory signal transduction through multiple pathways (Sakurai, 2012). TAK1 is involved in a wide variety of cellular processes, including cell survival, cell migration and invasion, inflammation, immune regulation, and tumorigenesis (Cho, Shim et al., 2021, Mukhopadhyay & Lee, 2020). Although the role of TAK1 has been widely studied, its precise role remains controversial. For example, by acting as a tumor suppressor, TAK1 has been shown to negatively associated with tumor progression in several human cancers including prostate cancer (Huang, Tang et al., 2021), hepatocellular carcinoma (HCC) (Tan, Zhao et al., 2020, Wang, Zhang et al., 2021), cervical cancer (Guan, Lu et al., 2017) and certain blood cancers (Guo, Zhang et al., 2019). In contrast, TAK1 promotes tumor progression in a variety of human cancers, including colon, ovarian, lung, and breast cancers (Augeri, Langenfeld et al., 2016, Cai, Shi et al., 2014, Xu, Niu et al., 2022). These aforementioned findings indicate that TAK1 plays a dual role in tumor initiation, progression, and metastasis. In our previous study, TAK1 has been shown to phosphorylate RASSF9 at serine 284 to inhibit cell proliferation by targeting the RAS/MEK/ERK axis in ESCC (Shi, Ju et al., 2021). To date, there have been no reports on the precise role of TAK1 in ESCC metastasis.

Phospholipase C epsilon 1 (PLCE1) is encoded by the *Plce1* gene on chromosome 10q23 in human, and belongs to the phospholipase C (PLC) family (Fukami, Inanobe et al., 2010, Kadamur & Ross, 2013). Like other PLC family members, PLCE1 is composed of a PLC catalytic domain, PH domain, EF domain, and C2 domain. In addition, PLCE1 contains unique regions, two C-terminal Ras association (RA) domains, and an N-terminal CDC25-homology domain (Kadamur & Ross, 2013). Once activated, PLCE1 is essential for intracellular signaling by catalyzing the hydrolysis of membrane phospholipids such as phosphatidylinositol 4,5-bisphosphate (PIP2), in order to produce two important secondary messengers, inositol 1,4,5-trisphosphate (IP3) and diacylglycerol (DAG), which further trigger IP3-dependent calcium ion (Ca^2+^) release from the endoplasmic reticulum and PKC activation (Fukami et al., 2010, Kadamur & Ross, 2013). Accumulating evidence has shown that PLCE1 promotes cell growth, migration, and metastasis in multiple human cancers, including hepatocellular carcinoma, non-small-cell lung cancer, head and neck cancer, bladder cancer, gastric cancer, and prostate cancer (Abnet, Freedman et al., 2010, Fan, Fan et al., 2019, Liao, Han et al., 2017, Ma, Wang et al., 2011, Ou, Guo et al., 2010, Wang, Liao et al., 2020, Yue, Zhao et al., 2019). However, whether and how PLCE1 affects cancer metastasis in ESCC remain largely unknown.

In the present study, we have examined the potential role of TAK1 in ESCC metastasis. We found that TAK1 negatively regulates cell migration and invasion in ESCC, and that PLCE1 is a downstream target of TAK1. TAK1 phosphorylates PLCE1 at serine 1060 (S1060) to inhibit its enzymatic activity, leading to decreased IP3 and DAG levels, both of which are products of PLCE1 catalyzed reactions. As a result, IP3/DAG triggered signal transduction in the axis of PKC/GSK-3β/β-Catenin was blunted. All these effects induced by TAK1 resulted in the repression of cell migration and invasion in ESCC.

## Results

### TAK1 negatively regulates ESCC migration and invasion

In our previous study, we showed that TAK1 expression was reduced in esophageal squamous tumor tissues, and that TAK1 inhibits ESCC proliferation (Shi et al., 2021). To examine whether TAK1 affects the epithelial-mesenchymal transition (EMT) process in ESCC, we increased TAK1 expression in ECA-109 cells by transfecting a plasmid expressing *Map3k7* (TAK1 gene name) and confirmed the overexpression of TAK1 (**Fig. 1A**). Owing to the promising role of epidermal growth factor (EGF) in EMT (Lu, Ghosh et al., 2003), it has been used to trigger EMT in ESCC. As shown in **Fig. 1B**, EGF treatment induced a spindle-shaped morphology in ECA-109 cells, which was markedly prevented by TAK1. These data implied that TAK1 is a negative regulator of EMT in ESCC. To address this hypothesis, we performed a transwell assay and found that TAK1 repressed the migration and invasion of ECA-109 cells (**Fig. 1C**; *SI Appendix,* Fig. S1A). The wound healing assay further confirmed the negative effects of TAK1 on cancer cell migration (**Fig. 1D**; *SI Appendix,* Fig. S1B). Next, we examined whether TAK1 affects EMT-related gene expression. Our data showed that TAK1 increased E-cadherin and ZO-1, two epithelial molecules, whereas mesenchymal molecules such as N-cadherin, Vimentin, and ZEB1 were reduced by TAK1 (**Fig. 1E**; *SI Appendix,* Fig. S1C). The qRT-PCR data confirmed these changes (*SI Appendix,* Fig. S1D).

**Figure 1.**
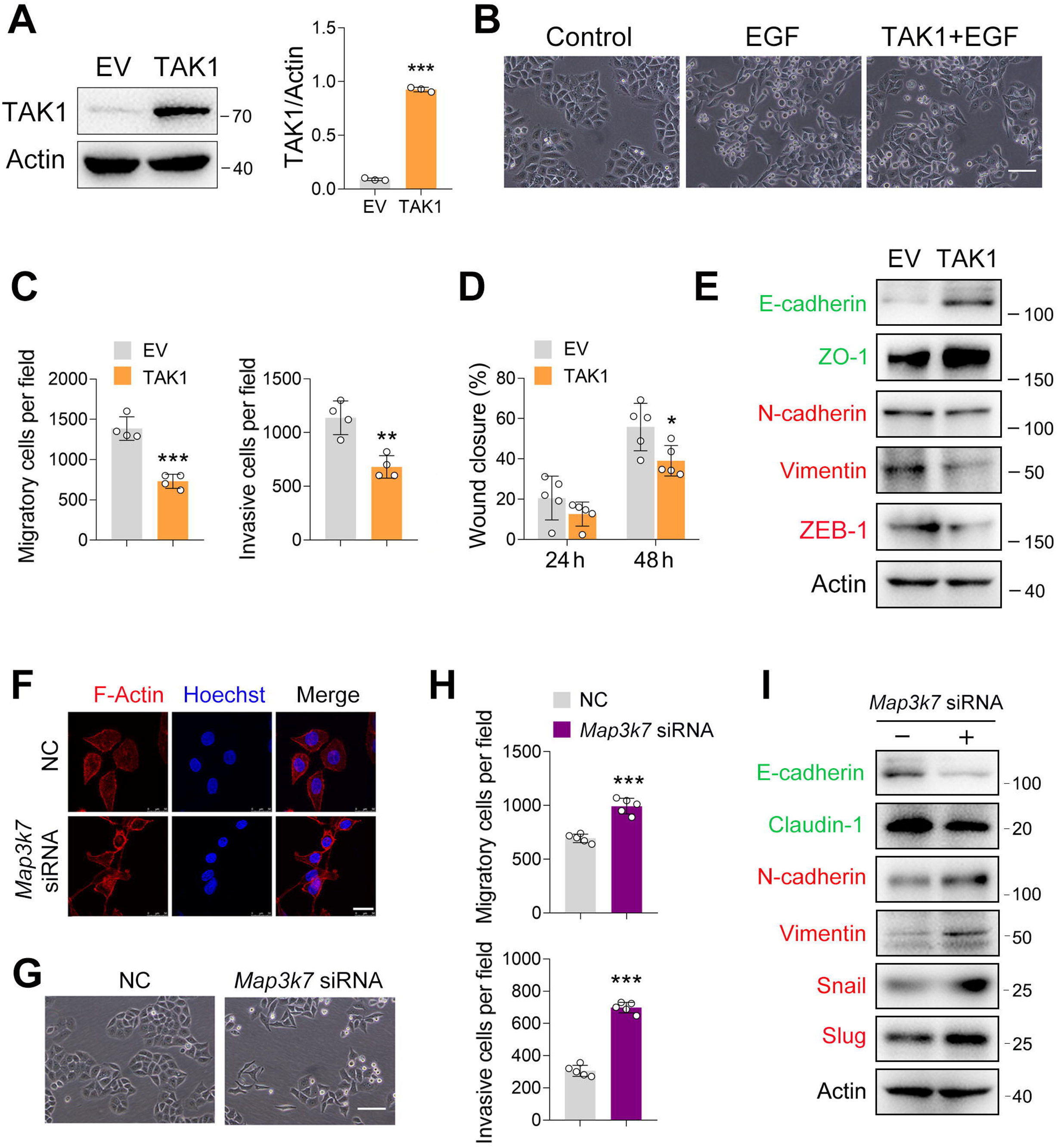
TAK1 negatively regultes ESCC migration and invasion. (A) Increased expression of TAK1 in ECA-109 cells transfected with a plasmid expressing *Map3k7*. (B) Increased expression of TAK1 inhibits the morphological changes to form spindle-shaped mesenchymal cells induced by EGF (100 ng/ml) in ECA-109. Scale bar = 100 µm. (C) Increased expression of TAK1 inhibits cell migration and invasion in ECA-109 cells. Cell migration and invasion were analyzed by transwell assay. *n* = 4 biologically independent replicates. (D) Wound healing assay showing cell migration was attenuated by TAK1. *n* = 5 biologically independent replicates. (E) TAK1 decreased mesenchymal marker gene expression, while increased the expression of epithelial markers. ECA-109 cells were transfected with the plasmid carrying *Map3k7*. 24 h post-transfection, protein samples were prepared and subjected to western blot. Actin was used as a loading control. (F) Knockdown of TAK1 increased the expression of F-Actin. ECA-109 cells were transfected with *Map3k7* siRNA. Seventy-two h post-transfection, cells were subjected to immunofluorescence analysis using an anti-F-Actin antibody (red). Hoechst was used to stain the nucleus (blue). Scale bar = 10 µm. (G) TAK1 knockdown induces spindle-shaped mesenchymal cell morphology in ECA-109 cells. Scale bar = 100 µm. (H) Reduced expression of TAK1 promotes cell migration and invasion. ECA-109 cells were transfected with *Map3k7* siRNA. 72 h post-transfection, cell migration and invasion were analyzed by transwell assay. *n* = 5 biologically independent replicates. (I) Knockdown of TAK1 increases mesenchymal protein marker expression, and decreases epithelial protein marker expression. Data are presented as mean ± SD. Statistical significance was tested by unpaired Student’s *t*-test. *\*p < 0.05*, *\*\*p < 0.01*, and *\*\*\*p < 0.001*.

Next, we performed loss-of-function experiments to examine the effect of TAK1 on ESCC cell migration and invasion. The knockdown efficiency of *Map3k7* siRNA on TAK1 expression has been reported previously (Shi et al., 2021). As EMT involves dynamic and spatial regulation of the cytoskeleton (Fife, McCarroll et al., 2014, Li & Wang, 2020), we analyzed the expression of F-Actin by immunofluorescence. As shown in **Fig. 1F**, TAK1 knockdown induced F-Actin expression. TAK1 knockdown promoted a spindle-shaped mesenchymal morphology in ECA-109 cells (**Fig. 1G**). In accordance with these changes, cell invasion and migration were stimulated, as evidenced by data from the wound healing and transwell assays (**Fig. 1H**; *SI Appendix,* Fig. S2A-B). The epithelial markers including E-cadherin and Claudin-1 were found to have decreased by TAK1 knockdown, whereas mesenchymal markers N-cadherin, Vimentin, Snail, and Slug were increased (**Fig. 1I**; *SI Appendix,* Fig. S2C). At the mRNA level, *Cdh1* and *Cldn1* were downregulated by TAK1 knockdown, whereas *Cdh2* and *Snail1* were upregulated (*SI Appendix,* Fig. S2D). To verify the effect of TAK1 knockdown on cell migration and invasion, we downregulated TAK1 expression using a lentivirus carrying *Map3k7* shRNA (LV-*Map3k7* shRNA) (*SI Appendix,* Fig. S3A, B). In LV-*Map3k7* shRNA-transduced cells, cell migration and invasion were clearly increased (*SI Appendix,* Fig. S3C-E). The epithelial markers E-cadherin, ZO-1, and Claudin-1 were repressed by TAK1 knockdown, whereas mesenchymal markers N-cadherin, Slug, and ZEB-1 were activated (*SI Appendix,* Fig. S3F, G). Our qRT-PCR data showed similar changes in these EMT genes (*SI Appendix,* Fig. S3H). Additionally, TAK1 knockdown was accomplished by CRISPR-Cas9 using *Map3k7* guide RNA (gRNA) (*SI Appendix,* Fig. S4A). Again, we observed that cell migration and invasion were enhanced by *Map3k7* gRNA (*SI Appendix,* Fig. S4B-D). The expression of E-cadherin, ZO-1, and Claudin-1 was inhibited by *Map3k7* gRNA, whereas the expression of N-cadherin, Vimentin, Snail, and Slug was activated (*SI Appendix,* Fig. S4E, F). Similar changes in these EMT-related gene expression were observed using qRT-PCR (*SI Appendix,* Fig. S4G). Furthermore, we inhibited TAK1 activity using Oxo, NG25, or Takinib and observed that all of these treatments induced morphological changes in ECA-109 cells from round shapes to spindle-like shapes (*SI Appendix,* Fig. S5A). Cell migration and invasion were also stimulated by TAK1 inhibition (*SI Appendix,* Fig. S5B-E). Taken together, these findings indicate that TAK1 negatively regulates ESCC migration and invasion.

### TAK1 phosphorylates PLCE1 at serine 1060

These aforementioned data clearly show that TAK1 negatively regulates the cell migration and invasion of ESCC. To reveal the underlying mechanism, we performed co-immunoprecipitation combined with mass spectrometry in order to identify potential downstream targets of TAK1. As previously reported, 24 proteins were phosphorylated in the immunocomplex (Shi et al., 2021). Of these, phospholipase C epsilon 1 (PLCE1) has attracted our attention because of its essential role in cell growth, migration, and metastasis in various human cancers (Abnet et al., 2010, Chen, Wang et al., 2019, Chen, Xin et al., 2020, Gu, Zheng et al., 2018, Kadamur & Ross, 2013, Wang Zhou et al., 2010). According to the mass spectrometry data, the serine residue at position 1060 (S1060) in PLCE1 was phosphorylated (**Fig. 2A**). Currently, there is no antibody against phosphorylated PLCE1 at S1060 (p-PLCE1 S1060), we therefore generated an antibody to detect p-PLCE1 S1060. To verify antibody specificity, S1060 in PLCE1 was mutated to alanine (S1060A). As shown in **Fig. 2B**, wild type (WT) PLCE1 was phosphorylated by TAK1 at S1060, whereas PCLE1 S1060A was resistant to TAK1 induced phosphorylation at S1060. Moreover, the p-PLCE1 S1060 induced by TAK1 was weakened by (5Z)-7-Oxozeaenol (Oxo), a potent inhibitor of TAK1 (**Fig. 2C**). Notably, phosphorylated TAK1 (p-TAK1) markedly decreased in the presence of Oxo, indicating that TAK1 was inhibited (**Fig. 2C**). To further confirm these findings, other TAK1 inhibitors, such as Takinib and NG25, were used, and similar changes in p-PLCE1 and p-TAK1 were observed (**Fig. 2D, E**). As for endogenous p-PLCE1 S1060, it was increased and decreased, respectively, by TAK1 overexpression and knockdown (*SI Appendix,* Fig. S6A, B). Moreover, the *in vitro* kinase assay showed that PLCE1 was phosphorylated by TAK1 at serine 1060 (*SI Appendix,* Fig. S6C). Collectively, these results indicate that PLCE1 is a downstream target of TAK1, and that PLCE1 S1060 is phosphorylated by TAK1. To further confirm this notion, we analyzed TAK1 and p-PCLE1 expression in clinical samples. As shown **Fig. 2F**, TAK1 expression was reduced in tumor tissues compared to that in their respective adjacent normal tissues. In accordance with these changes in TAK1, p-PLCE1 levels were also decreased in tumor tissues (**Fig. 2G**). Pearson’s correlation tests showed that TAK1 expression was positively correlated with p-PLCE1 expression (**Fig. 2H**). These changes in TAK1 and p-PLCE1 levels were confirmed by western blot data (**Fig. 2I**).

**Figure 2.**
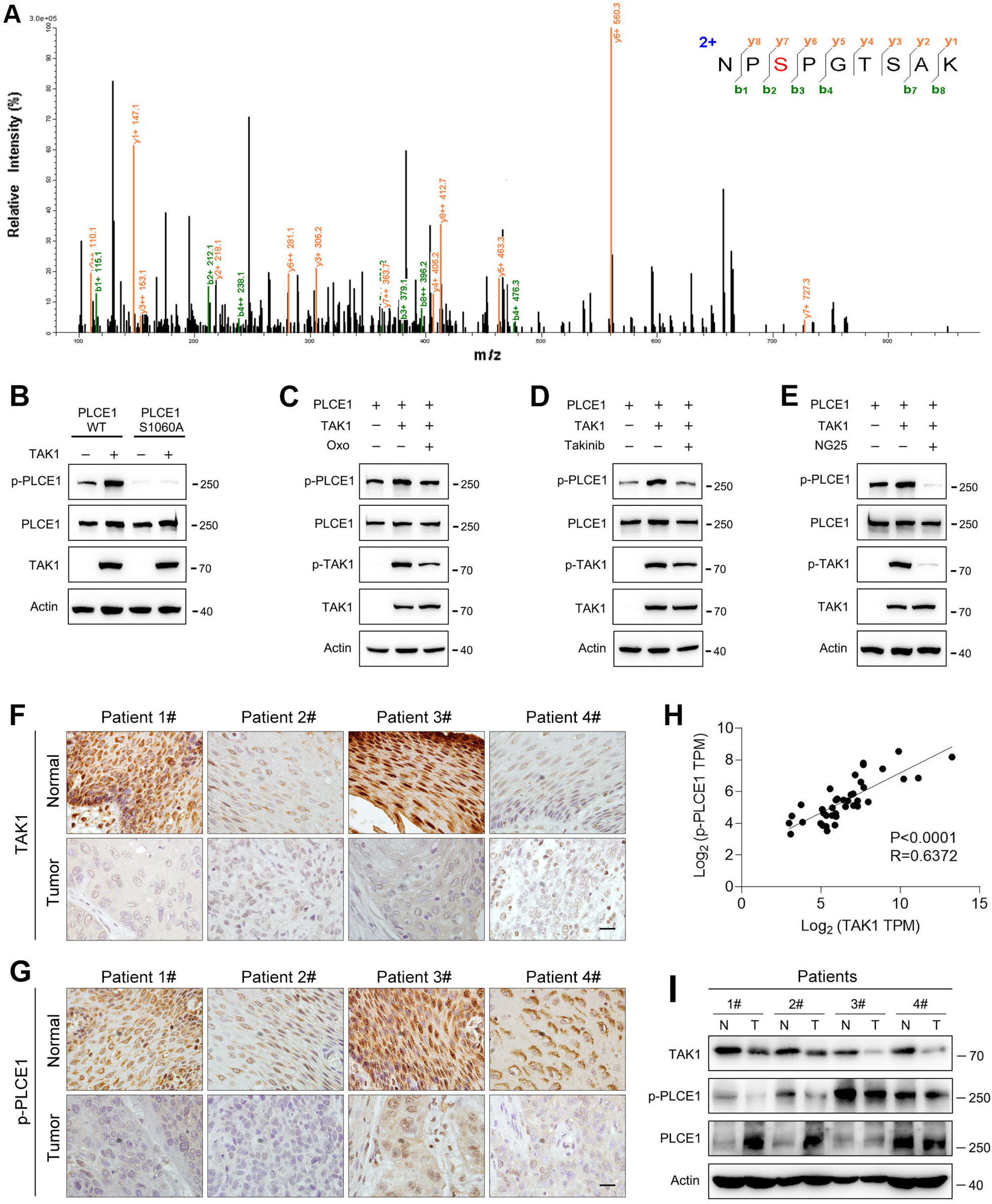
TAK1 phosphorylates PLCE1 at serine 1060. (A) Tandem mass spectrometry showing serine 1060 (S1060) in PLCE1 was phosphorylated by TAK1. ECA-109 cells were transfected with a plasmid expressing *Map3k7*. Twenty-four h-post transfection, cells were harvested and subjected to co-immunoprecipitation. The resulting immunocomplex was analyzed by liquid chromatography coupled with tandem mass spectrometry (LC-MS/MS). (B) TAK1 fails to phosphorylate PLCE1^S1060A^. ECA-109 cells were co-transfected with the plasmids carrying wildtype (WT) *Plce1*, mutated *Plce1* (PLCE1^S1060A^), or *Map3k7* as indicated. Twenty-four h-post transfection, cells were collected for western blot analysis (C-E) Inhibition of TAK1 reduces PLCE1 phosphorylation at S1060. ECA-109 cells were co-transfected with the plasmids expressing *Plce1* or *Map3k7*. 6 h post-transfection, TAK1 inhibitor (5Z)-7-Oxozeaenol (Oxo; 10 µM) (C), or 10 µM Takinib (D), or 10 µM NG25 (E) was added in culture medium, and cells were cultured for additional 18 h. Cells were then subjected to western blot analysis. Actin was used as a loading control (F-G) Immunohistochemical analysis of TAK1 (F) and p-PLCE1 (G) expression in normal and esophageal squamous tumor tissues. *n* = 4 biologically independent replicates. Scale bar = 20 µm. (H) Correlation between p-PLCE1 and TAK1 based on immunohistochemical data as shown in (F-G). 10 views for each sample were randomly chosen for Pearson correlation test. (I) TAK1 and p-PLCE1 protein levels in clinical samples. Protein levels were analyzed by western blot, and Actin was used as a loading control. *n* = 4 biologically independent replicates. N: normal tissue; T: tumor tissue.

### TAK1 phosphorylates PLCE1 to inhibit cell migration and invasion in ESCC

It has been well documented that PLCE1 plays a key role in cancer progression (Abnet et al., 2010, Chen et al., 2019, Chen et al., 2020, Gu et al., 2018, Wang et al., 2010). Therefore, we predicted that the negative effects of TAK1 on ESCC migration and invasion may rely on PLCE1. To verify this hypothesis, we first examined whether PLCE1 affects cell migration and invasion. In ECA-109 cells, PLCE1 overexpression significantly increased the migration and invasion (**Fig. 3A-C**; *SI Appendix*, Fig. S7A, B). In accordance with these changes, the epithelial marker E-cadherin was repressed by PLCE1, whereas the mesenchymal markers including Vimentin, Snail, Slug, and ZEB-1 were activated (**Fig. 3D**; *SI Appendix,* Fig. S7C). The qRT-PCR data also revealed that *Cdh1* was downregulated by PLCE1, while *Vim*, *Snail1*, and *Snail2* were upregulated (*SI Appendix,* Fig. S7D). To confirm these findings, we knocked down PLCE1 using siRNA. As shown in **Fig. 3E**, *Plce1* expression was markedly reduced by all three tested siRNAs. Similar changes were observed in the western blot data (**Fig. 3F**). Among these siRNAs, siRNA-2# was found to exhibit the highest knockdown efficiency and was selected for subsequent experiments. On the contrary, as compared to PLCE1 overexpression, PLCE1 knockdown impeded cell migration and invasion (**Fig. 3G, H**; *SI Appendix,* Fig. S8A, B). The expression patterns of genes involved in EMT were contrary to those of PLCE1 overexpression (**Fig. 3I**; *SI Appendix,* Fig. S8C, D). Moreover, similar to PLCE1 overexpression, the activation of PLCE1 by m-3M3FBS potentiated cell migration and invasion (*SI Appendix,* Fig. S9A-D). PLCE1 inhibition by U-73122 recapitulated the phenotypes induced by PLCE1 knockdown (*SI Appendix,* Fig. S10A-D).

**Figure 3.**
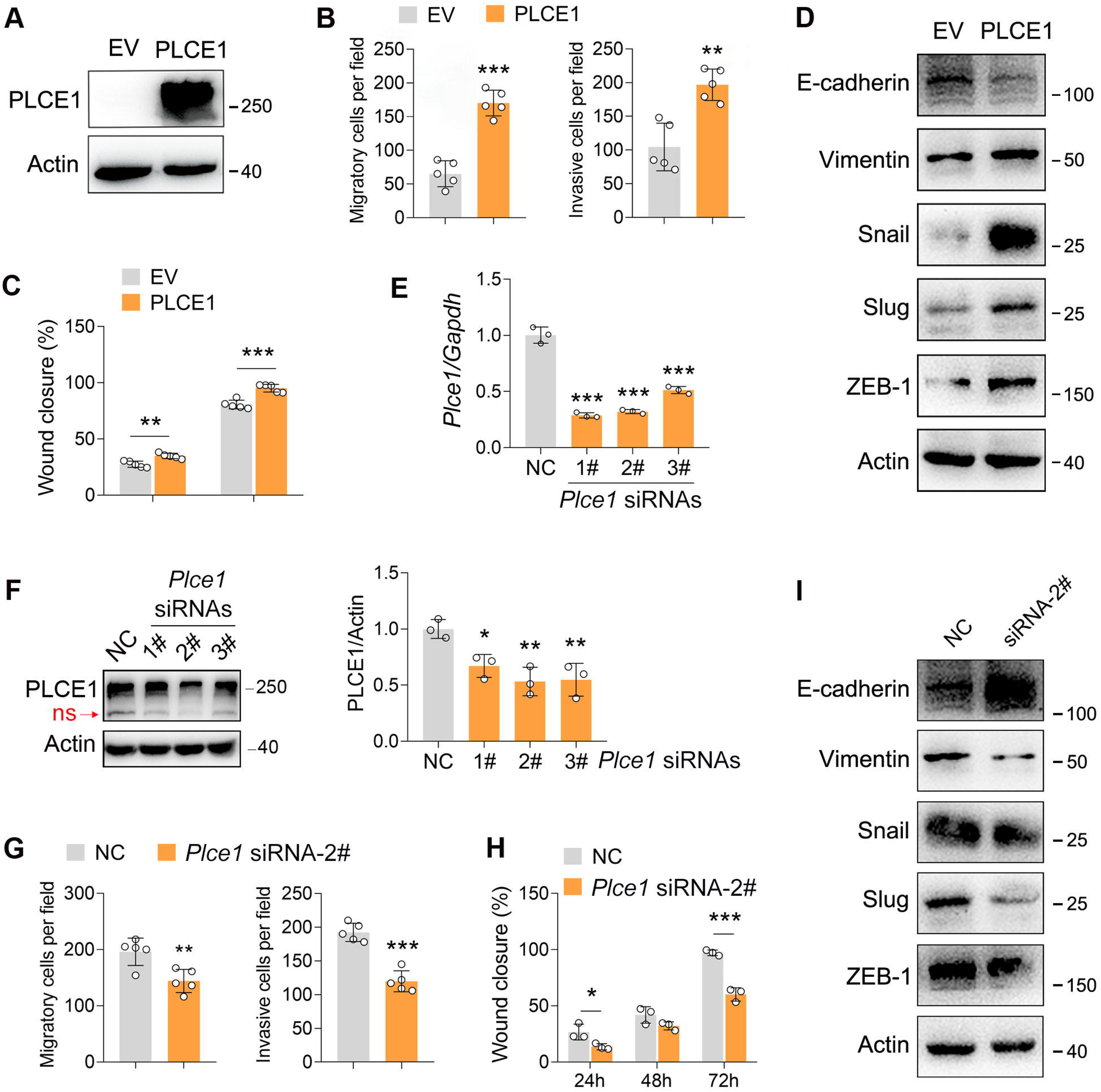
PLCE1 positively regulates ESCC migration and invasion. (A) Increased expression of PLCE1 in ECA-109 cells transfected with a plasmid expressing *Plce1*. (B-C) Increased expression of PLCE1 enhances cell migration and invasion. ECA-109 cells were transfected with the plasmid carrying *Plce1*. Twenty-four h-post transfection, cells were subjected to transwell (B) or wound healing (C) assay. *n* = 5 biologically independent replicates. (D) Increased expression of PLCE1 in ECA-109 cells induces mesenchymal protein marker expression, while reduces epithelial protein marker expression. (E-F) Knockdown of PLCE1. ECA-109 cells were transfected with siRNAs targeting *Plce1*. Seventy-two h-post transfection, cells were harvested for analyzing PLCE1 expression by qRT-PCR (E) and western blot (F). (G-H) Reduced expression of PLCE1 inhibits cell migration and invasion. ECA-109 cells were transfected with *Plce1* siRNA-2. Forty-eight h-post transfection, cells were subjected to transwell (G) or wound healing (H) assay. *n* = 3-5 biologically independent replicates. (I) Knockdown of PLCE1 promotes epithelial protein marker expression, while represses mesenchymal protein marker expression. ECA-109 cells were transfected with *Plce1* siRNA-2. Seventy-two h-post transfection, cells were harvested and subjected to western blot analysis. Protein level was detected by western blot, and Actin was used as a loading control. Gene expression was analyzed by qRT-PCR and *Gapdh* was used as a house-keeping gene. Data are presented as mean ± SD. Statistical significance was tested by unpaired Student’s *t*-test. *\*p < 0.05*, *\*\*p < 0.01*, and *\*\*\*p < 0.001*.

Since PLCE1 is a lipid hydrolase, we next investigated whether TAK1-induced phosphorylation of PLCE1 affects its enzymatic activity. To this end, we transfected ECA-109 cells with a plasmid bearing Myc-PLCE1 or TAK1, and PLCE1 was captured by pulldown using anti-Myc beads, which was then subjected to enzyme activity assay. As shown in **Fig. 4A**, the PLCE1 activity was reduced in the presence of TAK1; however, the PLCE1 S1060A activity was not affected (**Fig. 4B**), indicating TAK1 induced phosphorylation at S1060 repressed PLCE1 activity. As a lipid hydrolase, PLCE1 catalyzes PIP2 hydrolysis to produce IP3 and DAG, both of which are secondary messengers involved in diverse cellular processes (Kadamur & Ross, 2013). Therefore, we examined these two products to verify the inhibitory effect of TAK1 on PLCE1 expression. Our data showed that the productions of IP3 and DAG were increased by PLCE1, which was largely counteracted by TAK1 (**Fig. 4C-E**). As a messenger, IP3 induces the endoplasmic reticulum to release Ca^2+^ into the cytoplasm (Kadamur & Ross, 2013). Hence, we analyzed cytoplasmic Ca^2+^ levels using Fluo-4 AM staining. As expected, the signals for the cytoplasmic Ca^2+^ were increased by PLCE1 in ECA-109 cells, and this trend was attenuated by TAK1 (**Fig. 4F, G**). We also directly detected cytoplasmic Ca^2+^ using a fluorospectrophotometer and observed similar changes (**Fig. 4H**). Flow cytometry analysis also evidenced the increase in the cytoplasmic Ca^2+^ induced by PLCE1 was prevented by TAK1 (**Fig. 4I**). This evidence indicates that TAK1 inhibits PLCE1 activity, which is likely due to TAK1 induced phosphorylation of PLCE1 at S1060. In order to further verify this hypothesis, a series of functional experiments was performed. For instance, we observed that TAK1 reversed PLCE1-induced cell migration and invasion; however, TAK1 failed to affect PLCE1 S1060A-induced cell migration and invasion (*SI Appendix,* Fig. S11A-D). In addition to PLCE1, some other signaling pathways were also activated by TAK1. For example, TAK1 overexpression induced increases in p-IKK, p-JNK, p-p38 MAPK, and p-ERK in ECA-109 cells, whereas TAK1 knockdown reduced these protein levels (*SI Appendix,* Fig. S12A-D). At this regard, PLCE1 is likely a unique substrate of TAK1 for transducing its inhibitory effects on ESCC migration and invasion, although some other signaling pathways are also affected by TAK1. Overall, these data clearly indicate that PLCE1 positively regulates ESCC migration and invasion. By inducing phosphorylation at S1060, TAK1 inhibits PLCE1 enzyme activity, thereby counteracting PLCE1-induced ESCC migration and invasion. PLCE1 is a downstream target of TAK1 that inhibits ESCC migration and invasion.

**Figure 4.**
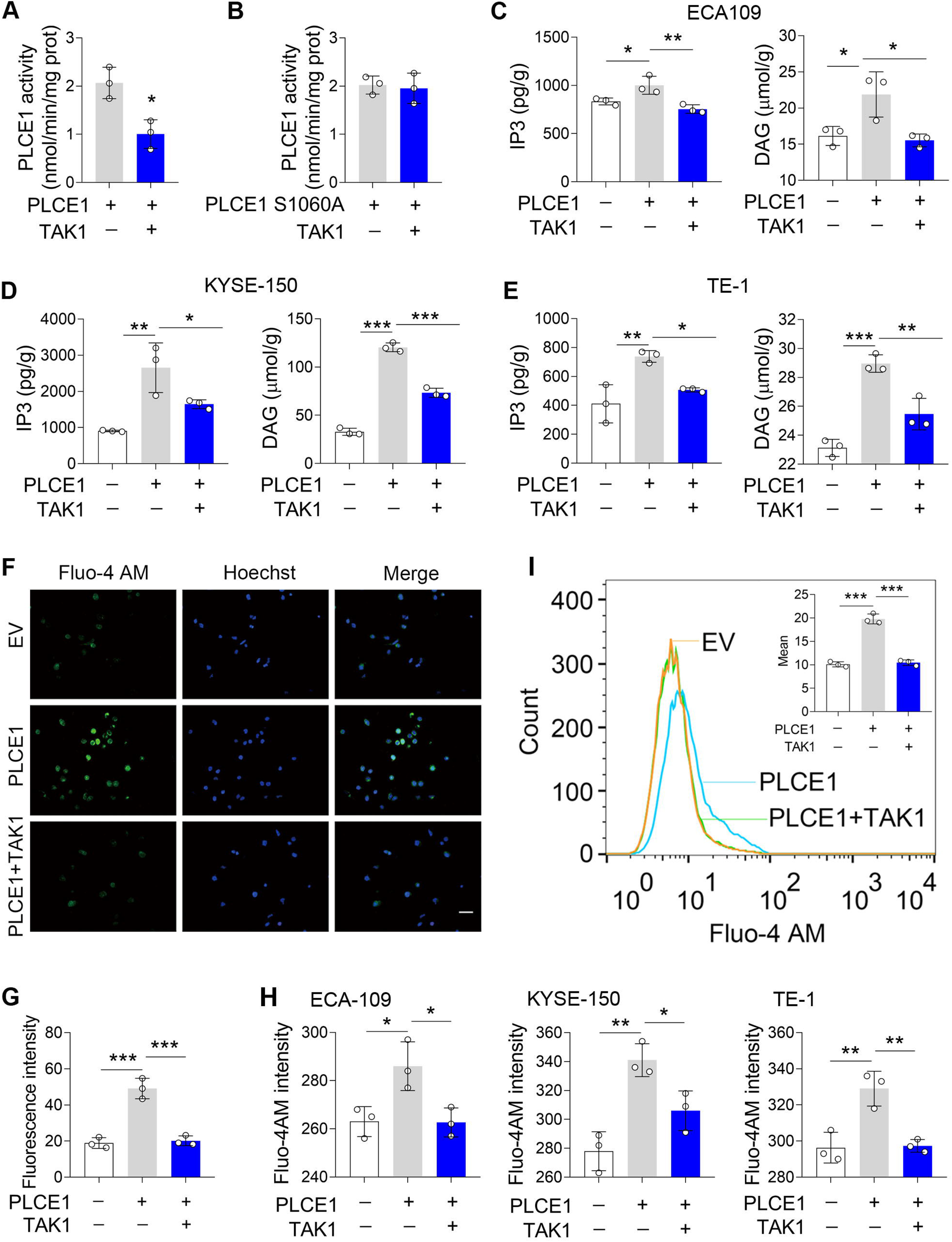
TAK1 inhibits PLCE1 enzyme activity. (A-B) Effects of TAK1 on PLCE1 (A) and PLCE1 S1060A (B) enzyme activity. ECA-109 cells were co-transfected with the plasmids expressing *Plce1-Myc* or *Map3k7*. Twenty-four h post-transfection, cells were subjected to pull down assay by using the beads with anti-Myc antibody. PLCE1 enzyme activity was assayed by Phospholipases C (PLC) Activity Assay Kit. *n* = 3 biologically independent replicates. (C-E) TAK1 abolishes PLCE1-induced IP3 and DAG in ECA-109 (C), KYSE-150 (D), and TE-1 cells (E). Cells were transfected with the plasmids bearing *Plce1* or *Map3k7* as indicated. Twenty-four h post-transfection, cells were harvested for measuring IP3 and DAG. *n* = 3 biologically independent replicates. (F) TAK1 attenuates PLCE1-induced intracellular Ca^2+^ ([Ca^2+^]). ECA-109 cells were transfected with the plasmids bearing *Plce1* or *Map3k7* as indicated. [Ca^2+^] was labeled with Fluo-4 AM, which was then detected by a fluorescent microscope. Scale bar = 20 µm. (G) Quantified fluorescence intensity of [Ca^2+^] in ECA-109 cells. *n* = 3 biologically independent replicates. (H) Fluorescence intensity of Fluo-4 in ECA-109, KYSE-150, and TE-1 cells was examined with a fluorospectrophotometer. *n* = 3 biologically independent replicates. (I) Flow cytometry analysis of [Ca^2+^]. Cell treatments were described in (F). Data are presented as mean ± SD. Statistical significance was tested by unpaired Student’s *t*-test (A-B) or two-tailed one-way ANOVA test (C-E, G-I). *\*p < 0.05*, *\*\*p < 0.01*, and *\*\*\*p < 0.001*.

### TAK1 inhibits PLCE1-induced signal transduction in the PKC/GSK-3β/β-Catenin axis

PLCE1 hydrolyzes PIP2 to produce two important secondary messengers, DAG and IP3, which trigger Ca^2+^ release from the endoplasmic reticulum into the cytoplasm to activate PKC (Harden, Hicks et al., 2009, Kadamur & Ross, 2013), suggesting that PKC activation is responsible for PLCE1-induced cell migration and invasion in ESCC. In order to test this hypothesis, we treated cells with 2-APB, an IP3 receptor (IP3R) inhibitor, and found that 2-APB treatment almost completely reversed PLCE1-induced the intracellular Ca^2+^ (*SI Appendix,* Fig. S13A-E). Accordingly, PLCE1-induced cell migration and invasion were replenished by 2-APB in ECA-109 cells (*SI Appendix,* Fig. S14A-D). The changes in EMT gene expression induced by PLCE1 were largely reversed by 2-APB treatment (*SI Appendix,* Fig. S14E). Similar to ECA-109 cells, 2-APB induced phenotypes in KYSE-150 and TE-1 cells (*SI Appendix,* Fig. S15A-E; Fig. S16A-E). BAPTA-AM, an intracellular Ca^2+^ chelator, dampened PLCE1-induced cell migration and invasion in ECA-109 cells (*SI Appendix,* Fig. S17A-D). PKC is a promising downstream target of intracellular Ca^2+^ (Kadamur & Ross, 2013). Therefore, we investigated whether the PLCE1-induced cell growth was dependent on PKC activation. To this end, midostaurin was used to inhibit PKC, and our data showed that midostaurin almost completely abolished cell migration and invasion induced by PLCE1 (*SI Appendix,* Fig. S18A-D).

It has been shown that PKC positively regulates the expression and stability of β-Catenin in a GSK-3β-dependent manner (Duong, Yu et al., 2017, Gwak, Cho et al., 2006, Liu, Shi et al., 2018, Ryu & Han, 2015, Tejeda-Munoz, Gonzalez-Aguilar et al., 2015). Of note, β-Catenin is considered as a positive regulator in the EMT process (Valenta, Hausmann et al., 2012). Therefore, we predicted that PLCE1 stimulates cell migration and invasion via the axis of PKC/GSK-3β/β-Catenin. To test this prediction, ECA-109 cells were treated with an IP3 receptor (IP3R) inhibitor (2-APB), Ca^2+^ chelator (BAPTA-AM), or PKC inhibitor (midostaurin), all of which successfully inhibited PKC activity, as evidenced by reduced phosphorylated PKC (p-PKC) (**Fig. 5A-C**; *SI Appendix,* S19A-C). As a result, phosphorylated GSK-3β (p-GSK3β) was decreased by 2-APB, BAPTA-AM, and Midostaurin (**Fig. 5A-C**; *SI Appendix,* S19A-C), indicating GSK-3β kinase activity was upregulated. Accordingly, phosphorylated β-Catenin (p-β-Catenin) was increased, leading to degradation of β-Catenin (**Fig. 5A-C**; *SI Appendix,* S19A-C). As a downstream target of β-Catenin, MMP2 expression was inhibited by all three tested chemicals (**Fig. 5A-C**; *SI Appendix,* S19A-C). The immunofluorescence data also showed that PLCE1-induced β-Catenin expression could be counteracted by 2-APB, BAPTA-AM, and Midostaurin (**Fig. 5D**). These results clearly indicate that PLCE1-induced cell migration and invasion in ESCC are likely via the axis of PKC/GSK-3β/β-Catenin. Furthermore, we observed that PLCE1-induced signal transduction in the axis of PKC/GSK-3β/β-Catenin could be inhibited by TAK1 (**Fig. 5E**; *SI Appendix,* S19D). The expression of β-Catenin induced by PLCE1 was largely reversed by TAK1 (**Fig. 5F**). However, inactive TAK1 (TAK1 K63W) showed no such effect (**Fig. 5G**; *SI Appendix,* S19E). Moreover, TAK1 failed to affect the PLCE1 S1060A induced signal cascade in the PKC/GSK-3β/β-Catenin axis (**Fig. 5H**; *SI Appendix,* S19F). This further indicates that TAK1 phosphorylates PLCE1 at S1060 to inhibit its activity and downstream signal transduction in the PKC/GSK-3β/β-Catenin axis.

**Figure 5.**
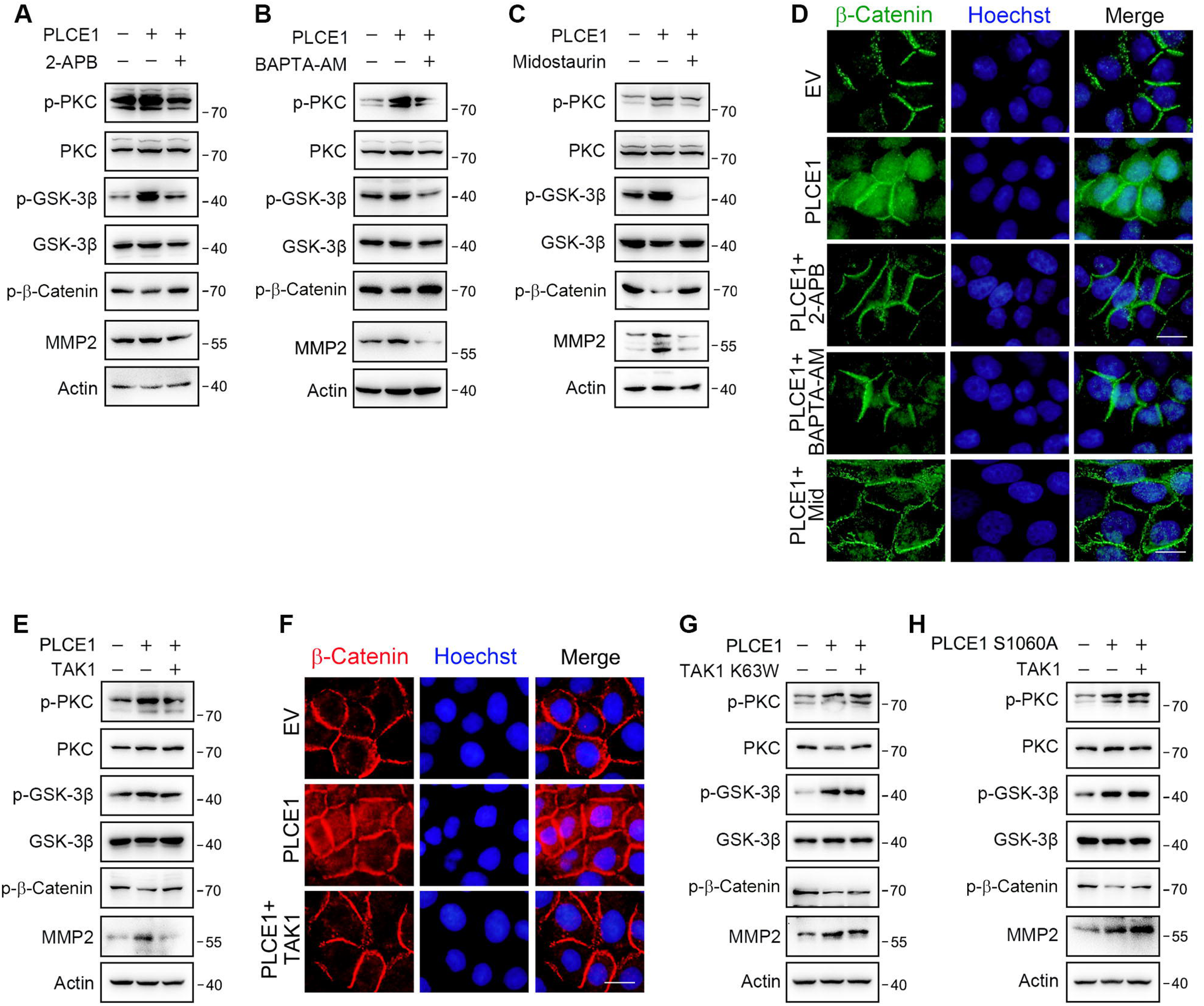
TAK1 inhibits PLCE1-induced signal transduction in the axis of PKC/GSK-3β/β-Catenin. (A) IP3R blocking inhibits PLCE1 induced signal transduction in the axis of PKC/GSK-3β/β-Catenin. ECA-109 cells were transfected with the plasmid expressing *Plce1* for 6 h and then treated with 2-APB (10 µM) for additional 18 h. (B) [Ca^2+^] blocking represses signal transduction in the axis of PKC/GSK-3β/β-Catenin induced by PLCE1. ECA-109 cells were transfected with the plasmid expressing *Plce1* for 6 h and then treated with BAPTA-AM (10 µM) for additional 18 h. (C) PKC inhibition blocks PLCE1 stimulated signal transduction in the axis of PKC/GSK-3β/β-Catenin. ECA-109 cells were transfected with the plasmid expressing *Plce1*. 6 h post-transfection, cells were treated with 100 nM of Midostaurin for additional 18 h. (D) PKC inhibition represses PLCE1-induced nuclear translocation of β-Catenin in ECA-109 cells. Cells were transfected with the plasmid expressing PLCE1. Six h post-transfection, 2-APB (10 µM), BAPTA-AM (10 µM), or Midostaurin (100 nM) was added in culture medium, and cells were cultured for additional 18 h. Scale bar = 10 µm. Immunofluorescence was used to examine subcellular distribution of β-Catenin. (E) TAK1 counteracts PLCE1-induced signal transduction in the axis of PKC/GSK-3β/β-Catenin. ECA-109 cells were transfected with the plasmids expressing *Plce1* or *Map3k7* as indicated for 24 h. (F) TAK1 reduces PLCE1-induced nuclear distribution of β-Catenin in ECA-109 cells. Cells were transfected with the plasmids expressing *Plce1* or *Map3k7* as indicated. Scale bar = 10 µm. (G) Dominant negative TAK1 (K63W) fails to block signal transduction in the axis of PKC/GSK-3β/β-Catenin/MMP2 induced by PLCE1. ECA-109 cells were transfected with the plasmids expressing *Plce1* or mutated *Map3k7* (TAK1 K63W) for 24 h. (H) TAK1 has no effect on PLCE1 S1060A induced signal transduction in the axis of PKC/GSK-3β/β-Catenin. ECA-109 cells were transfected with the plasmids expressing PLCE1 S1060A or TAK1 for 24 h. Protein levels were analyzed by western blot, and Actin was used as a loading control. Representative blots were shown.

### TAK1 inhibition promotes ESCC metastasis *in vivo*

To examine TAK1-regulated ESCC metastasis *in vivo*, a xenograft model with nude mice was used. Mice were injected with ECA109 cells via the tail vein, and treated daily with Takinib (50 mg/kg) or corn oil (vehicle) for consecutive 15 days. Eight weeks later, the mice were sacrificed for analysis of cancer cell metastasis. We found that four of six mice in the Takinib group developed lung metastasis, while only one of six mice in the vehicle group developed lung metastasis. Moreover, the number of metastatic nodules in lung tissues was higher in Takinib-treated mice (**Fig. 6A-C**). In addition, the effects of TAK1 inhibition on the axis of PKC/GSK-3β/β-Catenin was examined. As shown in **Fig. 6D**, TAK1 inhibition by Takinib activated PKC, leading to increased p-GSK-3β levels. As a result, p-β-Catenin was reduced and MMP2 expression was upregulated (**Fig. 6D, E**). Notably, the p-TAK1 levels were decreased by Takinib, indicating that TAK1 activity was successfully inhibited (**Fig. 6D, E**). Accordingly, p-PLCE1 expression was decreased in Takinib-treated mice (**Fig. 6D, E**). The body weight growth rate was not affected by Takinib (*SI Appendix,* Fig. S20A). These data indicate that TAK1 represses ESCC metastasis *in vivo*, and this benefit is likely due to TAK1-mediated phosphorylation of PLCE1 at S1060.

**Figure 6.**
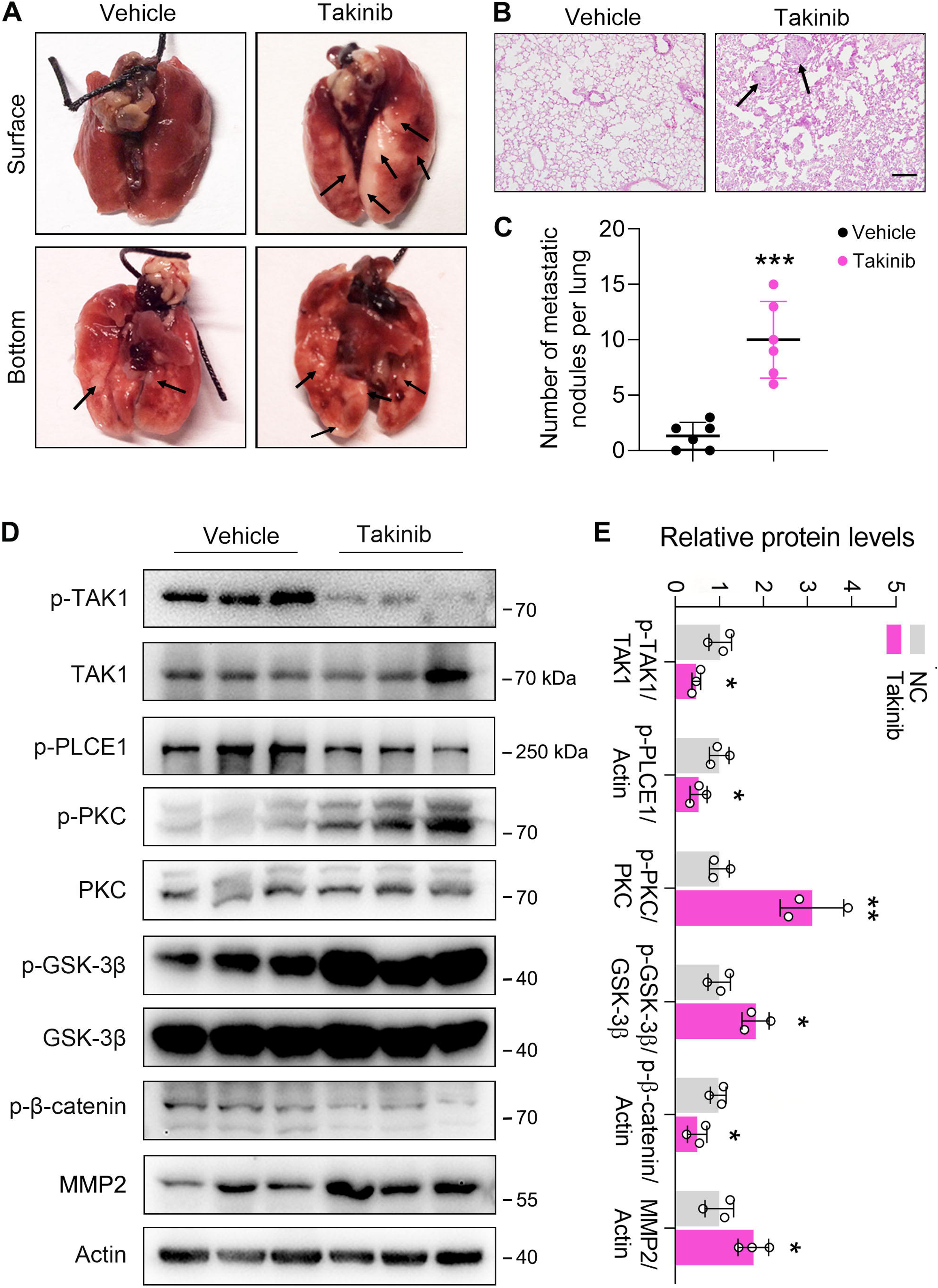
Inhibition of TAK1 by Takinib promotes ESCC metastasis in nude mice. Each mouse was intravenously injected with 1 x 10^6^ ECA109 cells diluted in 100 µl PBS. Mice were treated with Takinib at the dosage of 50 mg/kg/day for 15 days, mice in control group were received vehicle (corn oil). Eight weeks later, mice were sacrificed, and the lungs and livers from each group were collected and photographed. (A) Typical images of specimens. (B) Hematoxylin and eosin staining of metastatic nodules in lungs. (C) The number of nodules in lungs. (D) Takinib treatment induces signal transduction in the axis of PKC/GSK-3β/β-Catenin. Protein levels were analyzed by western blot, and Actin was used as a loading control. *n* = 3 biologically independent replicates. (E) Quantitative analysis of the western blot data shown in (D). Data are presented as mean ± SD. Statistical significance was tested by unpaired Student’s *t*-test. *\*p < 0.05, **p < 0.01*, and *\*\*\*p < 0.001*.

### PLCE1 facilitates ESCC metastasis *in vivo*

To further verify the function of PLCE1 in ESCC metastasis, we generated a stable cell line with PLCE1 knockdown using a lentivirus bearing PLCE1 shRNA (LV-shPLCE1) (*SI Appendix,* Fig. S20B-C). Cells with low PLCE1 expression were injected into nude mice via the tail vein to generate a mouse xenograft tumor model. Eight weeks later, the mice were sacrificed, and the nodule number and incidence rate were reduced in mice injected with LV-shPLCE1 cells (**Fig. 7A-C**). In addition, western blot analysis showed that PLCE1 silencing inhibited signal transduction in the PKC/GSK-3β/β-Catenin axis, as evidenced by reduced p-PKC and p-GSK-3β, and increased p-β-Catenin (**Fig. 7D, E**). Accordingly, MMP2 was decreased (**Fig. 7D, E**). These results further confirm that PLCE1 plays a key role in cancer cell metastasis in ESCC.

**Figure 7.**
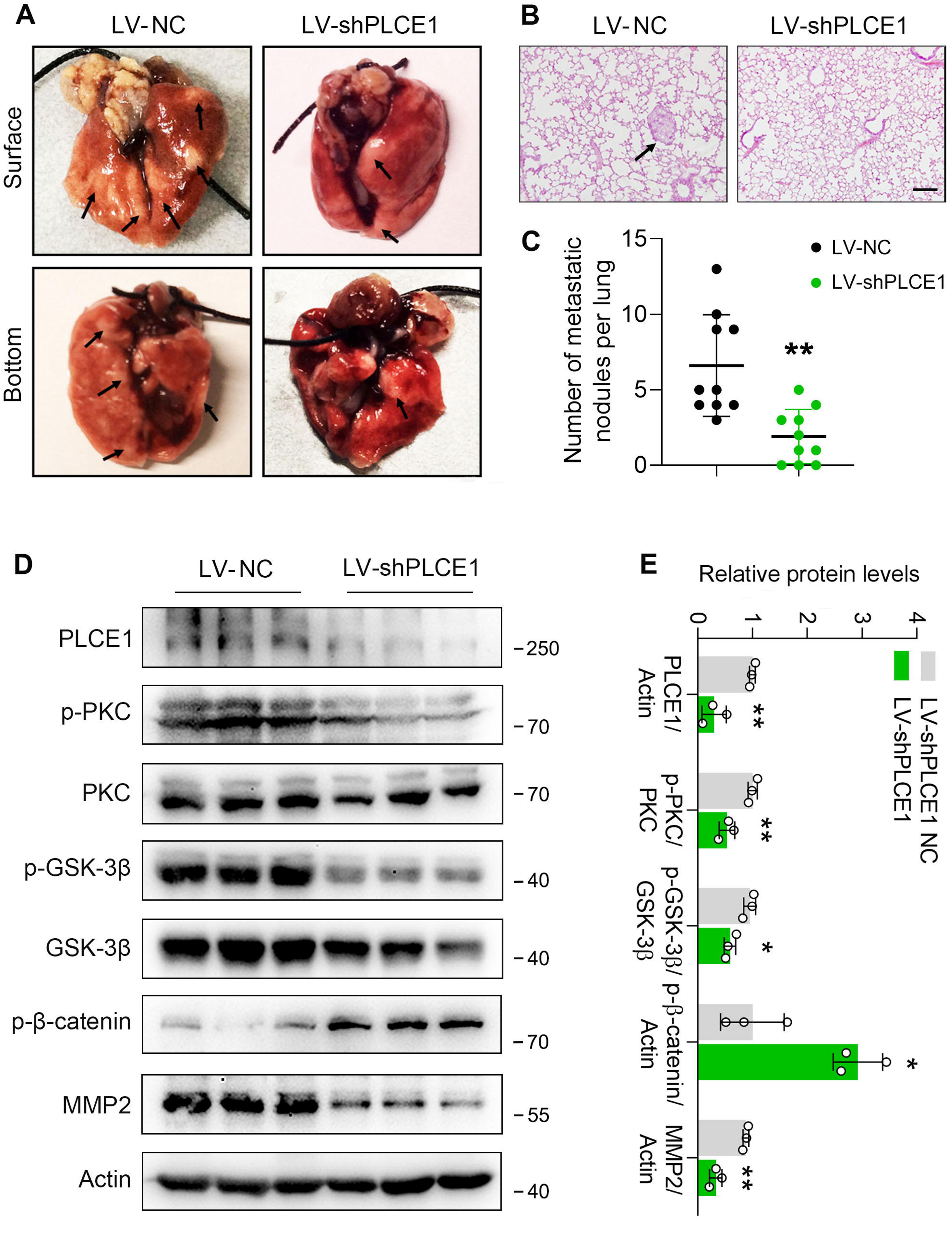
PLCE1 knockdown inhibits ESCC metastasis in nude mice. ECA-109 cells were transduced with lentivirus bearing PLCE1 shRNA (LV-shPLCE1) or NC shRNA (LV-shPLCE1 NC). Each mouse was intravenously injected with the LV transduced cells (1 x 10^6^ cells/mouse). Eight weeks later, mice were sacrificed, and the lungs and livers from each group were collected and photographed. (A) Typical images of lung specimens. (B) Hematoxylin and eosin staining of metastatic nodules in lungs. (C) The number of nodules in lungs. (D) PLCE1 knockdown represses signal transduction in the axis of PKC/GSK-3β/β-Catenin. Protein levels were analyzed by western blot, and Actin was used as a loading control. *n* = 3 biologically independent replicates. (E) Quantitative analysis of the western blot data shown in (D). Data are presented as mean ± SD. Statistical significance was tested by unpaired Student’s *t*-test. *\*\*p < 0.01*, and *\*\*\*p < 0.001*.

## Discussion

TAK1 is a serine/threonine kinase and a major member of the MAPK family (Zhu, Lama et al., 2021). In response to various cytokines, pathogens, lipopolysaccharides, hypoxia, and DNA damage, the E3 ligases TRAF2 and TRAF6 activate TAK1 through Lys63-linked polyubiquitylation, and then TAK1 undergoes auto-phosphorylation at Ser192 and Thr184/187 to achieve full activation (Skaug, Jiang et al., 2009, Sorrentino, Thakur et al., 2008). Consequently, activated TAK1, in turn, phosphorylates downstream substrates in order to initiate the NF-kB and MAPKs (i.e. ERK, p38 MAPK, and JNK) signaling pathways, thereby participates in cellular inflammation, immune response, fibrosis, cell death, and cancer cell invasion and metastasis (Yang, Chen et al., 2022, Zhou, Tao et al., 2021). TAK1 has been shown to decrease in high-grade human prostate cancer, and TAK1 deficiency promotes prostate tumorigenesis by increasing androgen receptor (AR) protein levels and activity or by activating the p38 MAPK pathway (Huang et al., 2021, Wu, Shi et al., 2012). Similarly, hepatocyte-specific TAK1 ablation drives RIPK1 kinase-dependent inflammation to promote liver fibrosis and hepatocellular carcinoma (Su, Gao et al., 2023, Tan et al., 2020, Xia, Ji et al., 2021). Furthermore, TAK1 represses the transcription of human telomerase and activates the tumor suppressor protein LKB1, indicating that TAK1 is a tumor suppressor (Adhikari, Xu et al., 2007, Fujiki, Miura et al., 2007, Xie, Zhang et al., 2006). Consistent with these findings, in our previous study, we found that TAK1 expression was reduced in esophageal squamous tumors when compared to that in adjacent normal tissues, and that TAK1 was negatively correlated with esophageal squamous tumor patient survival (Shi et al., 2021). In this study, we extended our research on TAK1 and observed that TAK1 inhibits cell migration and invasion in ESCC. In contrast, TAK1 plays a positive role in tumor cell proliferation, migration, invasion, colony formation, and metastasis, especially in breast, pancreatic, and non-small-cell lung cancers (Kim, Kim et al., 2023, Santoro, Zanotto et al., 2020, Tripathi, Shin et al., 2019). Therefore, TAK1 may play a pleiotropic role in various cancer cell types.

As a kinase, TAK1 is considered as a central regulator of tumor cell proliferation, migration and invasion and has been demonstrated to play tumor suppression or activation role via the NF-κB and MAPK activation (Mukhopadhyay & Lee, 2020, Wang et al., 2021, Zhang, Cheng et al., 2023). However, the precise molecular and cellular mechanisms through which TAK1 regulates ESCC metastasis remain unclear. As previously reported, using co-immunoprecipitation coupled with mass spectrometry (MS/MS), we identified RASSF9 as a downstream target of TAK1 to repress on cell proliferation in ESCC (Shi et al., 2021). By phosphorylating RASSF9 at S284, TAK1 negatively regulates ESCC proliferation (Shi et al., 2021). We also found that PLCE1 was present in the immunocomplex and was phosphorylated at S1060 in the presence of TAK1 in ECA-109 cells. It should be mentioned that, TAK1 inhibitors cannot completely abolish p-PLCE1 S1060 in cells and mice, which might be due to some other kinases also target PLCE1 at S1060. As a member of the human phosphoinositide-specific PLC family, PLCE1 is activated by various intracellular and extracellular signaling molecules, including hormones, cytokines, neurotransmitters, and growth factors (Kadamur & Ross, 2013). Numerous studies have shown that PLCE1 plays a key role in cancer development and progression via various pathways (Abnet et al., 2010, Fan et al., 2019, Ghosh, Nataraj et al., 2021, He, Wang et al., 2016, Yue et al., 2019). Moreover, PLCE1 has been identified as a susceptibility gene for ESCC (Abnet et al., 2010, Wang et al., 2010). This suggests that PLCE1 is a potential downstream target for transducing the negative effects of TAK1 on ESCC migration and invasion.

Once activated, PLCE1 catalyzes the hydrolysis of PIP2 on the cell membrane in order to produce two secondary messengers IP3 and DAG; IP3 induces Ca^2+^ release from the ER into the cytoplasm via IP3R; both Ca^2+^ and DAG are potent activators of PKC. PLCE1 regulates cell growth, proliferation, and differentiation, thereby playing a key role in tumor growth and development (Bunney & Katan, 2010, Kadamur & Ross, 2013, Land & Rubin, 2017, Smrcka, Brown et al., 2012). Consistent with these observations, we observed that PLCE1-overexpressing cells exhibited a higher production of intracellular IP3, DAG, and intracellular Ca^2+^. We found that PLCE1 overexpression accelerated cell migration and invasion, while PLCE1 knockdown resulted in the opposite phenotype. Moreover, in the presence of TAK1, PLCE1-induced cell migration and invasion were largely counteracted. Considering that TAK1 is a protein kinase, we propose that TAK1 phosphorylates PLCE1 at S1060 to inhibit its enzymatic activity, thus impedes cell migration and invasion in ESCC. Indeed, PLCE1 activity was blunted in the presence of TAK1, whereas the mutated PLCE1 (PLCE1 S1060A) was not affected. It has been shown that PKC positively regulates the expression and stability of β-Catenin in a GSK-3β-dependent manner (Duong et al., 2017, Gwak et al., 2006, Ryu & Han, 2015). Therefore, we investigated the impact of PLCE1 on the PKC/GSK-3β/β-Catenin axis. As expected, our data revealed that PLCE1 promotes cell migration and invasion in ESCC by activating the PKC/GSK-3β/β-Catenin pathway, results in a series of phosphorylation modification in PKC and GSK-3β. As a result, phosphorylated β-Catenin decreased, leading to a higher stability of β-Catenin, which eventually promotes the EMT process by affecting epithelial and mesenchymal gene expression.

In summary, our findings have indicated TAK1 negatively regulates the cell migration and invasion of ESCC. Mechanistically, TAK1 phosphorylates PLCE1 at residue S1060 to inhibit phospholipase activity, leading to a reduction in IP3 and DAG. Consequently, PKC activity was blunted, which results in decreased phosphorylated GSK-3β and increased phosphorylated β-Catenin. As a consequence, the stability of β-Catenin was decreased and its transcriptional activity was blunted. Thus, epithelial marker gene expression was upregulated by TAK1, whereas the mesenchymal marker gene expression was downregulated. These outcomes eventually lead to a breakdown in the cell migration and invasion of ESCC. Hence, our data revealed a new facet of TAK1 in the EMT process in ESCC by inhibiting PLCE1 activity and its downstream signal transduction in the axis of PKC/GSK-3β/β-Catenin. Moreover, TAK1 and, together with its downstream target, PLCE1, are potential drug targets for the development of agents for treating ESCC.

## Methods

### Cell culture

Human ESCC cell lines ECA109, KYSE-150 and TE-1 were purchased from the Shanghai Institute of Biochemistry and Cell Biology (Shanghai, China). HEK-293 cells were purchased from American Type Culture Collection (ATCC, Manassas, VA, USA). All cells were cultured in Dulbecco’s modified Eagle’s (DMEM) medium (Hyclone, UT, USA) containing 10% fetal bovine serum (FBS, Gibco, Carlsbad, CA), 100 U/ml penicillin and 100 μg/ml streptomycin (Life Technologies, Carlsbad, CA, USA) at 37°C in a humidified atmosphere of 5% CO_2_.

### Human esophageal squamous tumor specimens

Human esophageal squamous tumor specimens were obtained from the Affiliated Hospital of Nantong University. Human sample collection for research was conducted in accordance with the recognized ethical guideline of Declaration of Helsinki and approved by the Ethics Committee of Affiliated Hospital of Nantong University.

### Generation of antibodies against phospho-PLCE1 (S1060)

A rabbit polyclonal antibody that recognizing phospho-PLCE1 (S1060) was raised against the c-WSARNPS(p)PGTSAK peptide at Absin (Shanghai, China). Briefly, the phosphorylated polypeptide c-WSARNPS(p)PGTSAK, was synthesized as the target peptide. Two rabbits were then immunized with the target peptide antigen. The non-phosphorylated polypeptide c-WSARNPSPGTSAK was used as a control. The rabbits were sacrificed and blood was collected for further purification. The prepared antibody was verified and stored at −20°C until use.

### CRISPR-Cas9 mediated TAK1 deletion

*Map3k7* was deleted using the CRISPR-Cas9 method described previously (Shi et al., 2021). Briefly, we designed an integrated vector based on OriP/EBNA1 (epiCRISPR) bearing gRNA, Cas9, and puromycin-resistance genes. ECA-109 cells were transfected with plasmids using Lipofectamine 2000 (Invitrogen), according to the manufacturer’s instructions. The cells were cultured in a medium containing 2.5 μg/ml puromycin for 1 week. Surviving stable cells were used for further experiments. The gRNA sequences are listed in Table S1.

### Protein extraction and western blot analysis

Western blotting was performed as previously described (Tan, Krueger et al., 2022). Briefly, Total protein was extracted from the cells and tissues using protein lysis buffer supplemented with protease and phosphatase inhibitors (Roche Applied Science, Penzberg, Germany). Protein concentrations were assayed by the Pierce™ BCA Protein Assay Kit (Thermo Scientific, Waltham, MA, USA). Subsequently, equal amounts of protein were separated with 6% or 10% SDS-PAGE, and then protein were transferred onto 0.45 μm PVDF membranes. After blocking with 5% BSA in TBST for 1 h, membranes were incubated with primary antibodies at 4°C overnight. The membranes were then incubated with horseradish peroxidase-conjugated secondary antibodies for 1 h at room temperature. Protein signals were visualized using an enhanced chemiluminescence (ECL) reagent (Thermo Fisher Scientific, Waltham, MA, USA) and quantitatively analyzed using ImageJ (NIH). Actin was used as a loading control.

### Measurement of intracellular-free Ca^2+^

Cells were transfected with plasmid expressing PLCE1 and/or TAK1 for 6 h and then treated with BAPTA-AM (10 µM), 2-APB (10 µM), or Midostaurin (100 nM) for additional 18 h, respectively. Intracellular Ca^2+^ levels in the treated cells were measured using a Fluo-4 AM Assay Kit (Beyotime, Jiangsu, China). The cells were then washed with PBS, and incubated with 1 μM Fluo-4 AM in PBS at 37°C for 30 min. The cells were then washed with PBS three times and further incubated for 20 min to ensure that Fluo-4 AM was completely transformed into Fluo-4. Finally, Fluo-4 fluorescence intensity was detected using a confocal laser scanning fluorescence microscope (Olympus BX51, Tokyo, Japan), a fluorescence microplate (BioTek Instruments, Inc.), or flow cytometry (FACSCalibur, BD Bioscience, Franklin Lakes, NJ, USA) at excitation wavelength of 488 nm and an emission wavelength of 520 nm to determine the change in intracellular Ca^2+^ concentration.

### PLCE1 pulldown and phospholipase activity assay

Protein pull-down was performed using a previously reported method (Shi et al., 2021). HEK293 cells were transfected with plasmids expressing either PLCE1-Myc or TAK1. 24 h post-transfection, the cells were harvested for preparing total cell lysates, which was then subjected to protein pulldown assay using the Myc-Tag beads (Sepharose® Bead Conjugate; #55464; CST, Beverley, MA, USA). After 4 h of incubation on a rotator, the unbound proteins were removed using ice-cold wash buffer. Finally, phospholipase activity bound to the beads was detected using a PLC Activity Assay Kit (Jining Shiye, Shanghai, China) according to the manufacturer’s protocol. PLC catalyzes the hydrolysis of O-(4-nitrophenyl) choline (NPPC) to produce p-nitrophenol (PNP), which has an absorption maximum at 405 nm. The magnitude of the PLC enzyme activity was calculated by detecting the rate of increase in PNP at 405 nm.

### IP3 and DAG assays

IP3 and DAG levels in the cells were assayed using a Human Inositol 1,4,5-triphosphate enzyme-linked Immunosorbent Assay (ELISA) Kit and a Human Diacylglycerol commercial ELISA Kit (Mlbio, Shanghai, China), according to the manufacturer’s instructions. The absorbance was measured at 450 nm using a microplate reader.

### Co-immunoprecipitation and MS/MS spectrometry

Co-immunoprecipitation and MS/MS assays were performed as described previously (Shi et al., 2021). Briefly, ECA-109 cells were transfected with a plasmid expressing *Map3k7*. At 36 h post transfection, cells were harvested for co-immunoprecipitation using an antibody against TAK1. The resulting immunocomplex was subjected to liquid chromatography-tandem mass spectrometry (LC-MS/MS) (Shanghai Applied Protein Technology Co., Ltd.). MS/MS spectra were searched using the MASCOT engine (Matrix Science, London, UK; version 2.4) against UniProKB human (161584 total entries, downloaded 20180105).

### Mouse xenograft metastasis model

For *in vivo* metastasis assay, 4-week-old male BALB/c nude mice (18-20 g) were purchased from Shanghai Slake Laboratory Animal Co., Ltd. (Shanghai, China) and randomly divided into two groups (n = 6 - 10 per group). The mice were housed in air-filtered laminar flow cages (5 mice/cage) with a 12 h light cycle and adequate food and water. To examine the role of PLCE1 on metastasis, each mouse was intravenously (i.v.) injected with 1 × 10^6^ ECA109 cells diluted in 100 µl PBS, which were transduced with LV-shPLCE1 or negative control virus. Additionally, to examine TAK1 inhibition on metastasis, each mouse was intravenously (i.v.) injected with 1 x 10^6^ ECA109 cells. The mice were then intraperitoneally injected with Takinib (50 mg/kg/day) or vehicle (corn oil) for 15 days. The body weight of the mice was measured once per week. Eight weeks later, the mice were sacrificed, and the lungs and livers were collected for further analysis. All animal experiments were performed in accordance with the institutional ethical guidelines for animal care, and approved by the Animal Experimentation Ethics Committee of Nantong University (Approval ID: SYXK (SU) 2017-0046).

### Statistical analysis

Statistical significance of differences between groups were tested by using analysis of variance (ANOVA) or unpaired Student’s *t-*test. Data were presented as mean ± SD from at least three independent experiments. All statistical analyses were performed using GraphPad Prism version 8.0 (GraphPad Software, La Jolla, CA, USA). * p < 0.05 was accepted as statistical significance.

## Data availability

The dataset regarding MS-based protein phosphorylation modifications can be found in Figshare https://doi.org/10.6084/m9.figshare.25271140.

## Acknowledgements

We would like to thank Friedhelm Hildebrandt from University of Michigan Drive for providing the plasmid expressing PLCE1. This work was supported by the National Natural Science Foundation of China (No. 32271193); the Natural Science Foundation of Shanghai (No. 21ZR1449800); the Scientific Project from Shanghai Municipal Health Commission (202240015); the Postgraduate Research & Practice Innovation Program of Jiangsu Province (KYCX23_3411); and the Priority Academic Program Development (PAPD) of Jiangsu Higher Education Institutions.

## Author contributions

QJ performed all the experiments with assistance from WS, MZ, and JC. XL designed oligonucleotide sequences for qRT-PCR, shRNA, siRNA, and mutated PLCE1. LW provided advice throughout the development of the project. CS, TW, and HS developed the study concept and experiment design, supervised the study, and wrote the manuscript with input from all the authors.

## References

Abnet CC, Freedman ND, Hu N, Wang Z, Yu K, Shu XO, Yuan JM, Zheng W, Dawsey SM, Dong LM, Lee MP, Ding T, Qiao YL, Gao YT, Koh WP, Xiang YB, Tang ZZ, Fan JH, Wang C, Wheeler W et al. (2010) A shared susceptibility locus in PLCE1 at 10q23 for gastric adenocarcinoma and esophageal squamous cell carcinoma. Nat Genet 42: 764–7

Adhikari A, Xu M, Chen ZJ (2007) Ubiquitin-mediated activation of TAK1 and IKK. Oncogene 26: 3214–26

Augeri DJ, Langenfeld E, Castle M, Gilleran JA, Langenfeld J (2016) Inhibition of BMP and of TGFbeta receptors downregulates expression of XIAP and TAK1 leading to lung cancer cell death. Mol Cancer 15: 27

Bunney TD, Katan M (2010) Phosphoinositide signalling in cancer: beyond PI3K and PTEN. Nat Rev Cancer 10: 342–52

Cai PC, Shi L, Liu VW, Tang HW, Liu IJ, Leung TH, Chan KK, Yam JW, Yao KM, Ngan HY, Chan DW (2014) Elevated TAK1 augments tumor growth and metastatic capacities of ovarian cancer cells through activation of NF-kappaB signaling. Oncotarget 5: 7549–62

Chen Y, Wang D, Peng H, Chen X, Han X, Yu J, Wang W, Liang L, Liu Z, Zheng Y, Hu J, Yang L, Li J, Zhou H, Cui X, Li F (2019) Epigenetically upregulated oncoprotein PLCE1 drives esophageal carcinoma angiogenesis and proliferation via activating the PI-PLCepsilon-NF-kappaB signaling pathway and VEGF-C/ Bcl-2 expression. Mol Cancer 18: 1

Chen Y, Xin H, Peng H, Shi Q, Li M, Yu J, Tian Y, Han X, Chen X, Zheng Y, Li J, Yang Z, Yang L, Hu J, Huang X, Liu Z, Huang X, Zhou H, Cui X, Li F (2020) Hypomethylation-Linked Activation of PLCE1 Impedes Autophagy and Promotes Tumorigenesis through MDM2-Mediated Ubiquitination and Destabilization of p53. Cancer Res 80: 2175–2189

Cho SH, Shim HJ, Park MR, Choi JN, Akanda MR, Hwang JE, Bae WK, Lee KH, Sun EG, Chung IJ (2021) Lgals3bp suppresses colon inflammation and tumorigenesis through the downregulation of TAK1-NF-kappaB signaling. Cell Death Discov 7: 65

Duong M, Yu X, Teng B, Schroder P, Haller H, Eschenburg S, Schiffer M (2017) Protein kinase C Ill stabilizes beta-catenin and regulates its subcellular localization in podocytes. J Biol Chem 292: 12100–12110

Fan J, Fan Y, Wang X, Niu L, Duan L, Yang J, Li L, Gao Y, Wu X, Luo C (2019) PLCepsilon regulates prostate cancer mitochondrial oxidative metabolism and migration via upregulation of Twist1. J Exp Clin Cancer Res 38: 337

Fife CM, McCarroll JA, Kavallaris M (2014) Movers and shakers: cell cytoskeleton in cancer metastasis. Br J Pharmacol 171: 5507–23

Fujiki T, Miura T, Maura M, Shiraishi H, Nishimura S, Imada Y, Uehara N, Tashiro K, Shirahata S, Katakura Y (2007) TAK1 represses transcription of the human telomerase reverse transcriptase gene. Oncogene 26: 5258–66

Fukami K, Inanobe S, Kanemaru K, Nakamura Y (2010) Phospholipase C is a key enzyme regulating intracellular calcium and modulating the phosphoinositide balance. Prog Lipid Res 49: 429–37

Ghosh S, Nataraj NB, Noronha A, Patkar S, Sekar A, Mukherjee S, Winograd-Katz S, Kramarski L, Verma A, Lindzen M, Garcia DD, Green J, Eisenberg G, Gil-Henn H, Basu A, Lender Y, Weiss S, Oren M, Lotem M, Geiger B et al. (2021) PD-L1 recruits phospholipase C and enhances tumorigenicity of lung tumors harboring mutant forms of EGFR. Cell Rep 35: 109181

Gu D, Zheng R, Xin J, Li S, Chu H, Gong W, Qiang F, Zhang Z, Wang M, Du M, Chen J (2018) Evaluation of GWAS-Identified Genetic Variants for Gastric Cancer Survival. EBioMedicine 33: 82–87

Guan S, Lu J, Zhao Y, Woodfield SE, Zhang H, Xu X, Yu Y, Zhao J, Bieerkehazhi S, Liang H, Yang J, Zhang F, Sun S (2017) TAK1 inhibitor 5Z-7-oxozeaenol sensitizes cervical cancer to doxorubicin-induced apoptosis. Oncotarget 8: 33666–33675

Guo D, Zhang A, Huang J, Suo M, Zhong Y, Liang Y (2019) Suppression of HSP70 inhibits the development of acute lymphoblastic leukemia via TAK1/Egr-1. Biomed Pharmacother 119: 109399

Gwak J, Cho M, Gong SJ, Won J, Kim DE, Kim EY, Lee SS, Kim M, Kim TK, Shin JG, Oh S (2006) Protein-kinase-C-mediated beta-catenin phosphorylation negatively regulates the Wnt/beta-catenin pathway. J Cell Sci 119: 4702–9

Harden TK, Hicks SN, Sondek J (2009) Phospholipase C isozymes as effectors of Ras superfamily GTPases. J Lipid Res 50 Suppl: S243–8

He S, Xu J, Liu X, Zhen Y (2021) Advances and challenges in the treatment of esophageal cancer. Acta Pharm Sin B 11: 3379–3392

He Y, Wang C, Wang Z, Zhou Z (2016) Genetic variant PLCE1 rs2274223 and gastric cancer: more to be explored? Gut 65: 359–60

Huang Z, Tang B, Yang Y, Yang Z, Shi L, Bai Y, Yan B, Karnes RJ, Zhang J, Jimenez R, Wang L, Wei Q, Yang J, Xu W, Jia Z, Huang H (2021) MAP3K7-IKK Inflammatory Signaling Modulates AR Protein Degradation and Prostate Cancer Progression. Cancer Res 81: 4471–4484

Kadamur G, Ross EM (2013) Mammalian phospholipase C. Annu Rev Physiol 75: 127–54

Kim MJ, Kim JY, Shin JH, Kang Y, Lee JS, Son J, Jeong SK, Kim D, Kim DH, Chun E, Lee KY (2023) FFAR2 antagonizes TLR2- and TLR3-induced lung cancer progression via the inhibition of AMPK-TAK1 signaling axis for the activation of NF-kappaB. Cell Biosci 13: 102

Lagergren J, Smyth E, Cunningham D, Lagergren P (2017) Oesophageal cancer. Lancet 390: 2383–2396

Land M, Rubin CS (2017) A Calcium- and Diacylglycerol-Stimulated Protein Kinase C (PKC), Caenorhabditis elegans PKC-2, Links Thermal Signals to Learned Behavior by Acting in Sensory Neurons and Intestinal Cells. Mol Cell Biol 37

Li X, Wang J (2020) Mechanical tumor microenvironment and transduction: cytoskeleton mediates cancer cell invasion and metastasis. Int J Biol Sci 16: 2014–2028

Liao X, Han C, Qin W, Liu X, Yu L, Zhu G, Yu T, Lu S, Su H, Liu Z, Chen Z, Yang C, Huang K, Liu Z, Liang Y, Huang J, Dong J, Li L, Qin X, Ye X et al. (2017) PLCE1 polymorphisms and expression combined with serum AFP level predicts survival of HBV-related hepatocellular carcinoma patients after hepatectomy. Oncotarget 8: 29202–29219

Liu T, Shi Y, Chan MTV, Peng G, Zhang Q, Sun X, Zhu Z, Xie Y, Sham KWY, Li J, Liu X, Ho IHT, Gin T, Lu Z, Wu WKK, Cheng CHK (2018) Developmental protein kinase C hyper-activation results in microcephaly and behavioral abnormalities in zebrafish. Transl Psychiatry 8: 232

Lu Z, Ghosh S, Wang Z, Hunter T (2003) Downregulation of caveolin-1 function by EGF leads to the loss of E-cadherin, increased transcriptional activity of beta-catenin, and enhanced tumor cell invasion. Cancer Cell 4: 499–515

Ma H, Wang LE, Liu Z, Sturgis EM, Wei Q (2011) Association between novel PLCE1 variants identified in published esophageal cancer genome-wide association studies and risk of squamous cell carcinoma of the head and neck. BMC Cancer 11: 258

Mao C, Zeng X, Zhang C, Yang Y, Xiao X, Luan S, Zhang Y, Yuan Y (2021) Mechanisms of Pharmaceutical Therapy and Drug Resistance in Esophageal Cancer. Front Cell Dev Biol 9: 612451

Mukhopadhyay H, Lee NY (2020) Multifaceted roles of TAK1 signaling in cancer. Oncogene 39: 1402–1413

Ou L, Guo Y, Luo C, Wu X, Zhao Y, Cai X (2010) RNA interference suppressing PLCE1 gene expression decreases invasive power of human bladder cancer T24 cell line. Cancer Genet Cytogenet 200: 110–9

Pennathur A, Gibson MK, Jobe BA, Luketich JD (2013) Oesophageal carcinoma. Lancet 381: 400–12

Ryu JM, Han HJ (2015) Autotaxin-LPA axis regulates hMSC migration by adherent junction disruption and cytoskeletal rearrangement via LPAR1/3-dependent PKC/GSK3beta/beta-catenin and PKC/Rho GTPase pathways. Stem Cells 33: 819–32

Sakurai H (2012) Targeting of TAK1 in inflammatory disorders and cancer. Trends Pharmacol Sci 33: 522–30

Santoro R, Zanotto M, Simionato F, Zecchetto C, Merz V, Cavallini C, Piro G, Sabbadini F, Boschi F, Scarpa A, Melisi D (2020) Modulating TAK1 Expression Inhibits YAP and TAZ Oncogenic Functions in Pancreatic Cancer. Mol Cancer Ther 19: 247–257

Shi H, Ju Q, Mao Y, Wang Y, Ding J, Liu X, Tang X, Sun C (2021) TAK1 Phosphorylates RASSF9 and Inhibits Esophageal Squamous Tumor Cell Proliferation by Targeting the RAS/MEK/ERK Axis. Adv Sci (Weinh) 8: 2001575

Skaug B, Jiang X, Chen ZJ (2009) The role of ubiquitin in NF-kappaB regulatory pathways. Annu Rev Biochem 78: 769–96

Smrcka AV, Brown JH, Holz GG (2012) Role of phospholipase Cepsilon in physiological phosphoinositide signaling networks. Cell Signal 24: 1333–43

Sorrentino A, Thakur N, Grimsby S, Marcusson A, von Bulow V, Schuster N, Zhang S, Heldin CH, Landstrom M (2008) The type I TGF-beta receptor engages TRAF6 to activate TAK1 in a receptor kinase-independent manner. Nat Cell Biol 10: 1199–207

Su W, Gao W, Zhang R, Wang Q, Li L, Bu Q, Xu Z, Liu Z, Wang M, Zhu Y, Wu G, Zhou H, Wang X, Lu L (2023) TAK1 deficiency promotes liver injury and tumorigenesis via ferroptosis and macrophage cGAS-STING signalling. JHEP Rep 5: 100695

Sung H, Ferlay J, Siegel RL, Laversanne M, Soerjomataram I, Jemal A, Bray F (2021) Global Cancer Statistics 2020: GLOBOCAN Estimates of Incidence and Mortality Worldwide for 36 Cancers in 185 Countries. CA Cancer J Clin 71: 209–249

Tan R, Krueger RK, Gramelspacher MJ, Zhou X, Xiao Y, Ke A, Hou Z, Zhang Y (2022) Cas11 enables genome engineering in human cells with compact CRISPR-Cas3 systems. Mol Cell 82: 852–867 e5

Tan S, Zhao J, Sun Z, Cao S, Niu K, Zhong Y, Wang H, Shi L, Pan H, Hu J, Qian L, Liu N, Yuan J (2020) Hepatocyte-specific TAK1 deficiency drives RIPK1 kinase-dependent inflammation to promote liver fibrosis and hepatocellular carcinoma. Proc Natl Acad Sci U S A 117: 14231–14242

Tejeda-Munoz N, Gonzalez-Aguilar H, Santoyo-Ramos P, Castaneda-Patlan MC, Robles-Flores M (2015) Glycogen Synthase Kinase 3beta Is Positively Regulated by Protein Kinase Czeta-Mediated Phosphorylation Induced by Wnt Agonists. Mol Cell Biol 36: 731–41

Tripathi V, Shin JH, Stuelten CH, Zhang YE (2019) TGF-beta-induced alternative splicing of TAK1 promotes EMT and drug resistance. Oncogene 38: 3185–3200

Valenta T, Hausmann G, Basler K (2012) The many faces and functions of beta-catenin. EMBO J 31: 2714–36

Wang L, Zhang X, Lin ZB, Yang PJ, Xu H, Duan JL, Ruan B, Song P, Liu JJ, Yue ZS, Fang ZQ, Hu H, Liu Z, Huang XL, Yang L, Tian S, Tao KS, Han H, Dou KF (2021) Tripartite motif 16 ameliorates nonalcoholic steatohepatitis by promoting the degradation of phospho-TAK1. Cell Metab 33: 1372–1388 e7

Wang LD, Zhou FY, Li XM, Sun LD, Song X, Jin Y, Li JM, Kong GQ, Qi H, Cui J, Zhang LQ, Yang JZ, Li JL, Li XC, Ren JL, Liu ZC, Gao WJ, Yuan L, Wei W, Zhang YR et al. (2010) Genome-wide association study of esophageal squamous cell carcinoma in Chinese subjects identifies susceptibility loci at PLCE1 and C20orf54. Nat Genet 42: 759–63

Wang XK, Liao XW, Yang CK, Liu ZQ, Han QF, Zhou X, Zhang LB, Deng T, Gong YZ, Huang JL, Huang R, Han CY, Yu TD, Su H, Ye XP, Peng T, Zhu GZ (2020) Oncogene PLCE1 may be a diagnostic biomarker and prognostic biomarker by influencing cell cycle, proliferation, migration, and invasion ability in hepatocellular carcinoma cell lines. J Cell Physiol 235: 7003–7017

Wu M, Shi L, Cimic A, Romero L, Sui G, Lees CJ, Cline JM, Seals DF, Sirintrapun JS, McCoy TP, Liu W, Kim JW, Hawkins GA, Peehl DM, Xu J, Cramer SD (2012) Suppression of Tak1 promotes prostate tumorigenesis. Cancer Res 72: 2833–43

Xia S, Ji L, Tao L, Pan Y, Lin Z, Wan Z, Pan H, Zhao J, Cai L, Xu J, Cai X (2021) TAK1 Is a Novel Target in Hepatocellular Carcinoma and Contributes to Sorafenib Resistance. Cell Mol Gastroenterol Hepatol 12: 1121–1143

Xie M, Zhang D, Dyck JR, Li Y, Zhang H, Morishima M, Mann DL, Taffet GE, Baldini A, Khoury DS, Schneider MD (2006) A pivotal role for endogenous TGF-beta-activated kinase-1 in the LKB1/AMP-activated protein kinase energy-sensor pathway. Proc Natl Acad Sci U S A 103: 17378–83

Xu G, Niu L, Wang Y, Yang G, Zhu X, Yao Y, Zhao G, Wang S, Li H (2022) HDAC6-dependent deacetylation of TAK1 enhances sIL-6R release to promote macrophage M2 polarization in colon cancer. Cell Death Dis 13: 888

Yang L, Chen H, Hu Q, Liu L, Yuan Y, Zhang C, Tang J, Shen X (2022) Eupalinolide B attenuates lipopolysaccharide-induced acute lung injury through inhibition of NF-kappaB and MAPKs signaling by targeting TAK1 protein. Int Immunopharmacol 111: 109148

Yue QY, Zhao W, Tan Y, Deng XL, Zhang YH (2019) PLCE1 inhibits apoptosis of non-small cell lung cancer via promoting PTEN methylation. Eur Rev Med Pharmacol Sci 23: 6211–6216

Zhang X, Cheng L, Gao C, Chen J, Liao S, Zheng Y, Xu L, He J, Wang D, Fang Z, Zhang J, Yan M, Luan Y, Chen S, Chen L, Xia X, Deng C, Chen G, Li W, Liu Z et al. (2023) Androgen Signaling Contributes to Sex Differences in Cancer by Inhibiting NF-kappaB Activation in T Cells and Suppressing Antitumor Immunity. Cancer Res 83: 906–921

Zhou Y, Tao T, Liu G, Gao X, Gao Y, Zhuang Z, Lu Y, Wang H, Li W, Wu L, Zhang D, Hang C (2021) TRAF3 mediates neuronal apoptosis in early brain injury following subarachnoid hemorrhage via targeting TAK1-dependent MAPKs and NF-kappaB pathways. Cell Death Dis 12: 10

Zhu L, Lama S, Tu L, Dusting GJ, Wang JH, Liu GS (2021) TAK1 signaling is a potential therapeutic target for pathological angiogenesis. Angiogenesis 24: 453–470

