## Supplemental Figures S1-S20 and supplemental Tables S1-S2 for "TAK1-mediated phosphorylation of PLCE1 represses PIP2 hydrolysis to impede esophageal squamous cancer metastasis"

**This PDF file includes the following:**

Figs. S1 to S20

Tables S1 and S2

Materials and methods

References

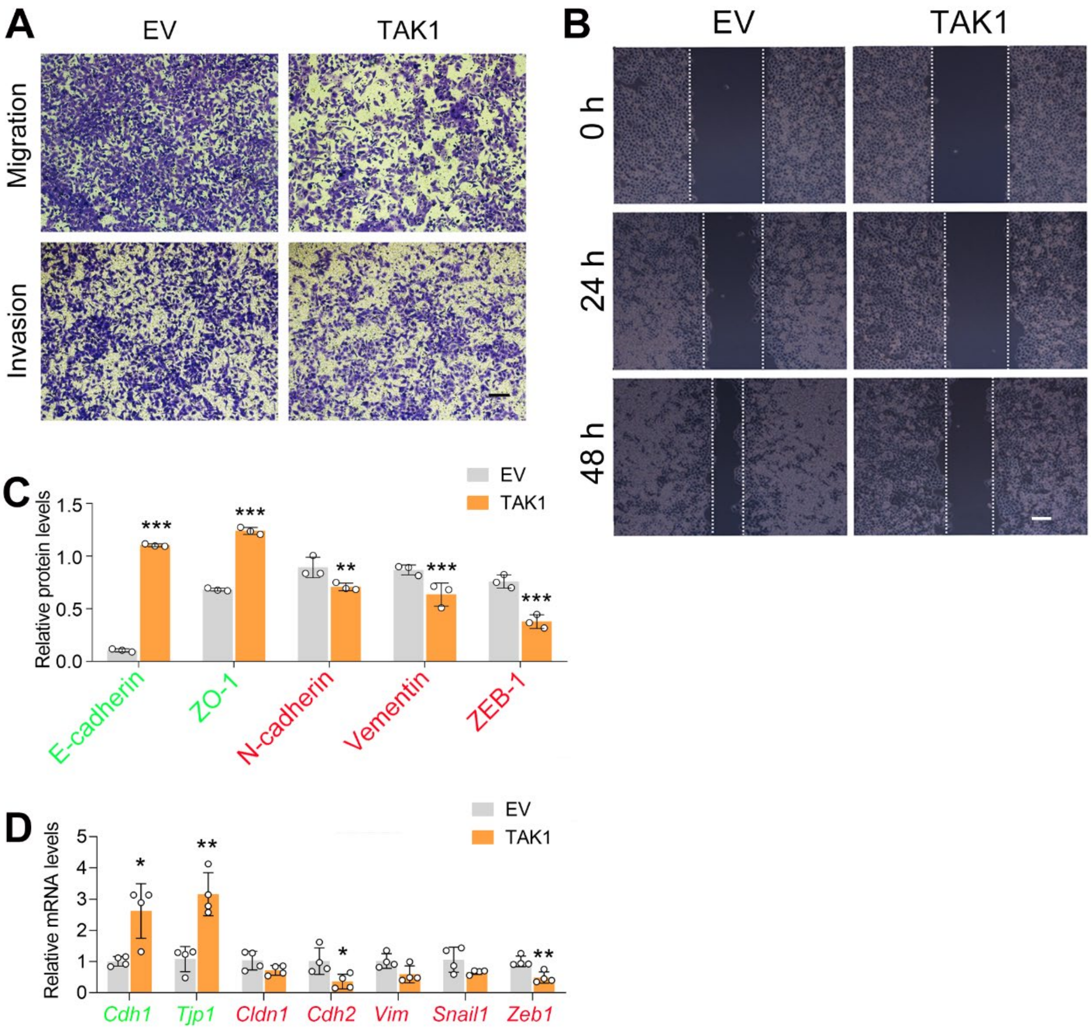


***Figure S1. TAK1 represses cell migration in ECA-109 cells.*** *Cells were transfected with a plasmid expressing Map3k7. 24 or 48 h-post transfection, cells were subjected to transwell (A) and wound healing assay (B).* Scale bar = 500 μm (A) and 100 μm (B). *C) Quantified protein expression based on western blot data in Fig. 1E. n* = 3 biologically independent replicates. *D,* TAK1 decreased mesenchymal marker gene expression, while increased the expression of epithelial markers. Gene expression was analyzed by qRT-PCR, and *Gapdh* was used as a house-keeping gene. *n* = 4 biologically independent replicates. Data are presented as mean ± SD. Statistical significance was tested by unpaired Student's *t*-test. **p < 0.05, **p < 0.01,* and ****p < 0.001*.

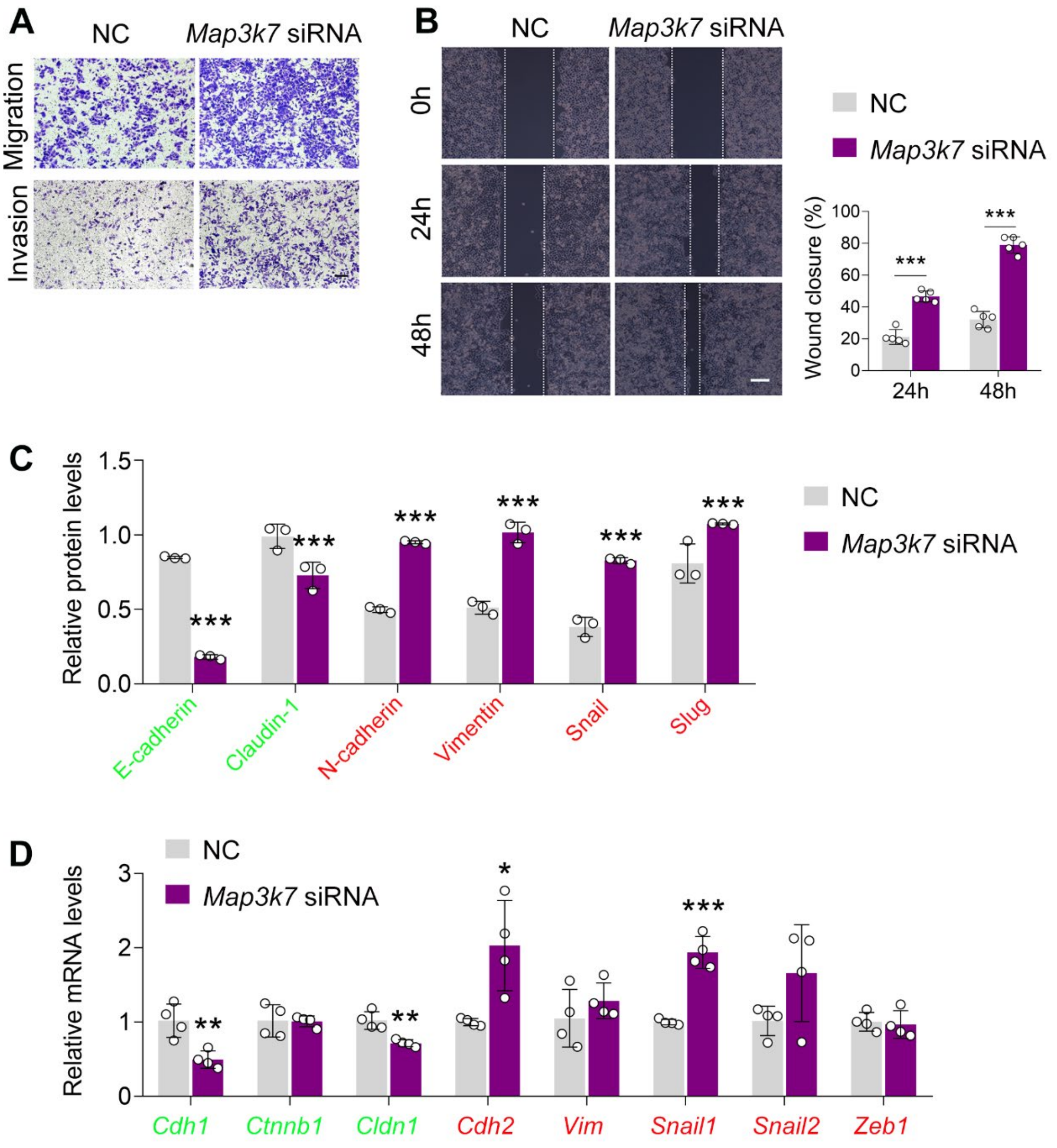


***Figure S2.*** **TAK1 silencing promotes ESCC migration and invasion.** A-C) Knockdown of TAK1 in ECA-109 cells promotes cell migration and invasion analyzed by transwell (A) and wound healing (B) assays. Scale bar = 500 μm (A) or 100 μm (B). C) Q*uantitative analysis of the western blot data as shown in Fig. 1I. n = 3 biologically independent replicates. D*) Reduced expression of TAK1 in ECA-109 cells increases mesenchymal marker gene expression, while decreases epithelial marker gene expression. Gene expression was analyzed by qRT-PCR, *Gapdh* was used as a house-keeping gene. *n* = 4 biologically independent replicates. Data are presented as mean ± SD. Statistical significance was tested by unpaired Student's *t*-test. **p < 0.05, **p < 0.01,* and ****p < 0.001*.

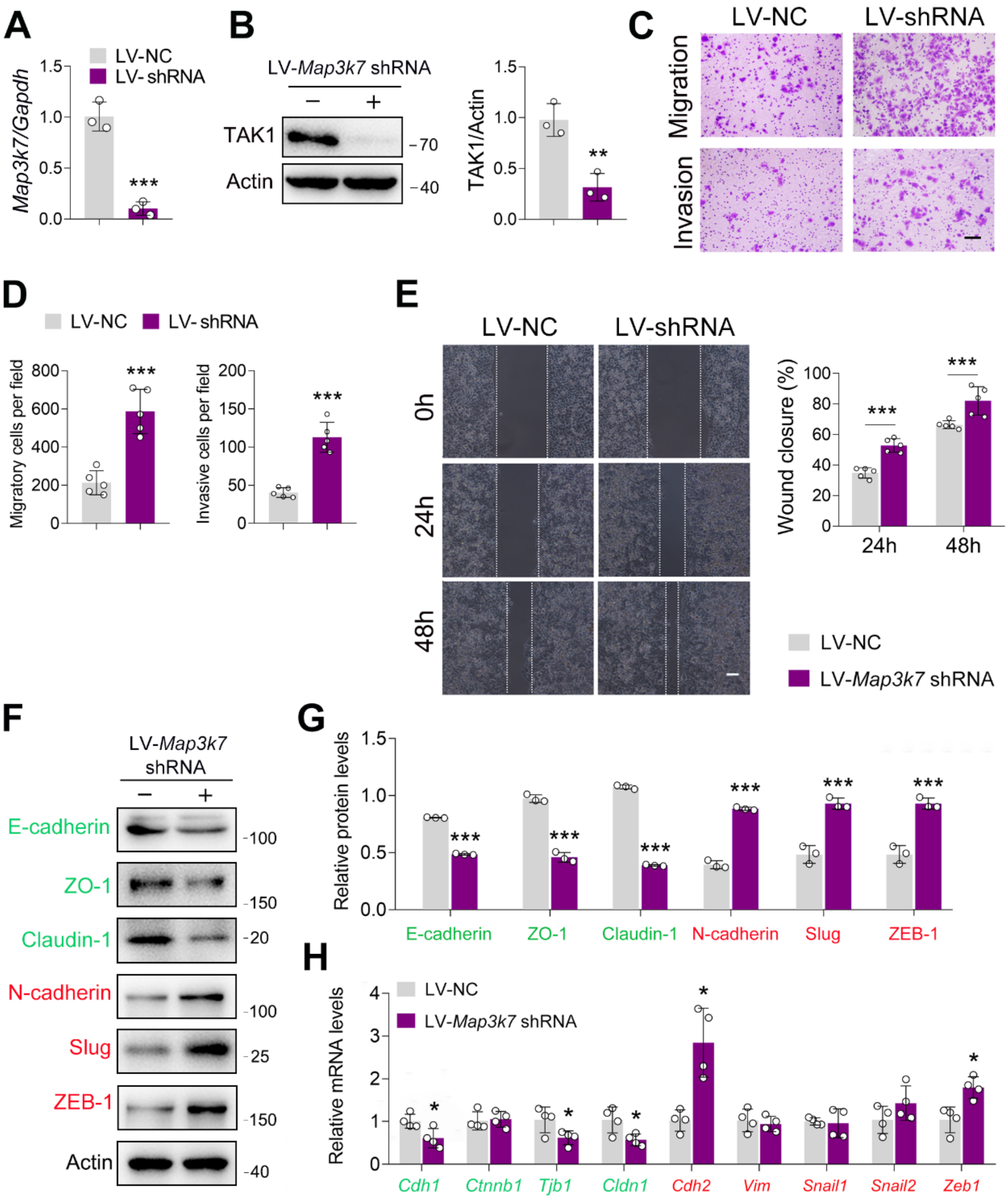


***Figure S3.*** **TAK1 knockdown facilitates ESCC migration and invasion.** A-B) Knockdown of TAK1 by *Map3k7* shRNA. ECA-109 cells were transduced with lentivirus bearing *Map3k7* shRNA (LV-*Map3k7* shRNA) or NC shRNA (LV-NC). Forty-eight h-post transduction, cells were harvested for analyzing TAK1 expression by qRT-PCR (A) and western blot (B). $n$ = 3 biologically independent replicates. C-E) *Decreased* expression of TAK1 facilitates cell migration and invasion. ECA-109 cells were transduced with LV-*Map3k7* shRNA or LV-NC. Forty-eight h-post transduction, cells were subjected to transwell (C, D) or wound healing (E)

assay. $n = 5$ biologically independent replicates. Scale bar = 500 μm (C); Scale bar = 100 μm (E). F-G) Reduced expression of TAK1 increases mesenchymal marker expression, and decreases epithelial marker expression. *n = 3 biologically independent replicates. H)* Reduced expression of TAK1 in ECA-109 cells affects EMT related gene expression. Protein levels were analyzed by western blot, and Actin was used as a loading control. Gene expression was detected by qRT-PCR, *Gapdh* was used as a house-keeping gene. $n = 4$ biologically independent replicates. Data are presented as mean ± SD. Statistical significance was tested by unpaired Student's *t*-test. *$p < 0.05$, **$p < 0.01$, and ***$p < 0.001$.

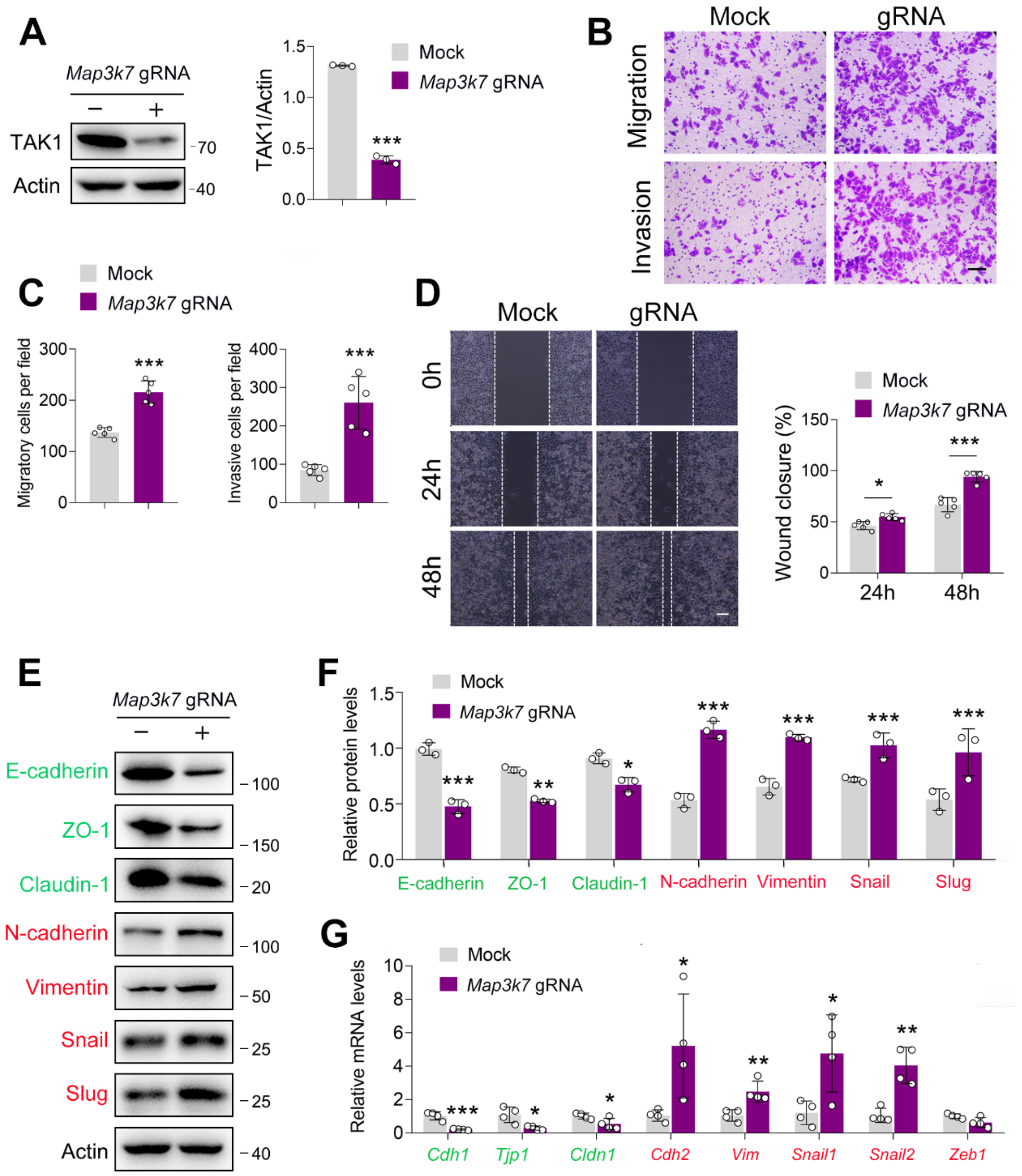


***Figure S4.*** **TAK1 knockout accelerates ESCC migration and invasion.** A-B) TAK1 expression was decreased by *Map3k7* gRNA in ECA-109 cells. TAK1 knockout was achieved by CRISPR-Cas9. B-D, Knockout of TAK1 expression in ECA-109 cells accelerates cell migration and invasion as analyzed by transwell (B, C) and wound healing (D) assays. Scale bar = 500 μm (B); Scale bar = 100 μm (D). E-F) Loss of TAK1 increases mesenchymal protein marker expression, and reduces epithelial protein marker expression. ECA-109 cells were treated with *Map3k7* gRNA, and then cells were harvested for western blot analysis. *n = 3 biologically independent replicates. G*) Knockout of TAK1 expression in ECA-109 cells alters EMT related gene expression as analyzed by qRT-PCR. *Gapdh* was used as a house-keeping gene. $n = 4$ biologically independent replicates. Protein levels were analyzed by western blot, and Actin was used as a loading control. Data are presented as mean ± SD. Statistical significance was tested by unpaired Student's *t*-test. *$p < 0.05$, **$p < 0.01$, and ***$p < 0.001$.

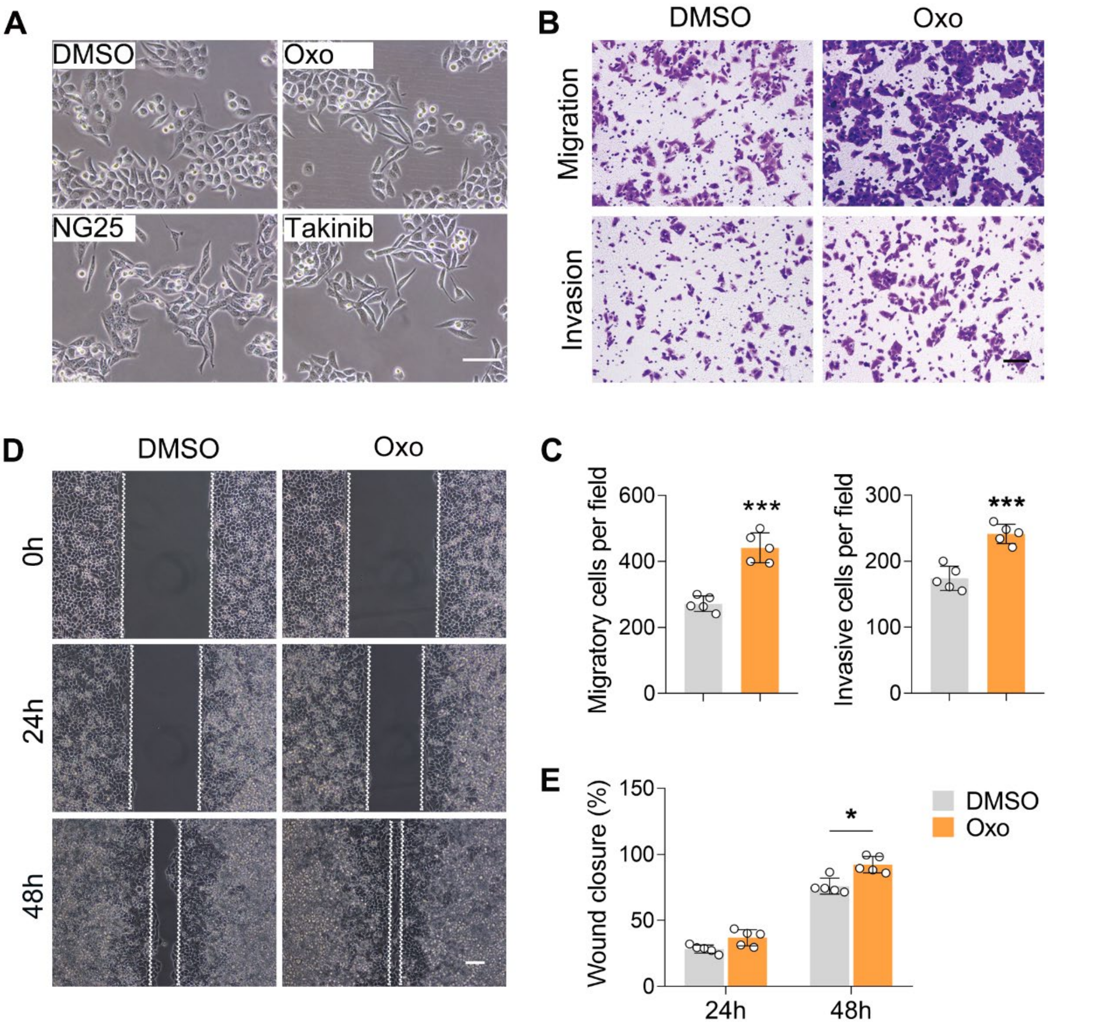


***Figure S5.*** **Inhibition of TAK1 potentiates cell migration and invasion in ECA-109 cells.** Cells were treated with (5Z)-7-Oxozeaenol (Oxo; 10 μM), or NG25 (10 μM), or Takinib (10 μM) for 24 h, and then cell morphology, migration and invasion were analyzed. A) Inhibition of TAK1 promotes morphological changes to form spindle-shaped mesenchymal cells in ECA-109 cells. Scale bar = 100 μm. B-E) Inhibition TAK1 promotes cell migration and invasion in ECA-109 cells. Cell migration and invasion were analyzed by transwell (B, C) and wound healing (D, E) assays. Scale bar = 500 μm (B); Scale bar = 100 μm (D). *n* = 5 biologically independent replicates. Data are presented as mean ± SD. Statistical significance was tested by unpaired Student's *t*-test. *$p < 0.05$, and ***$p < 0.001$.

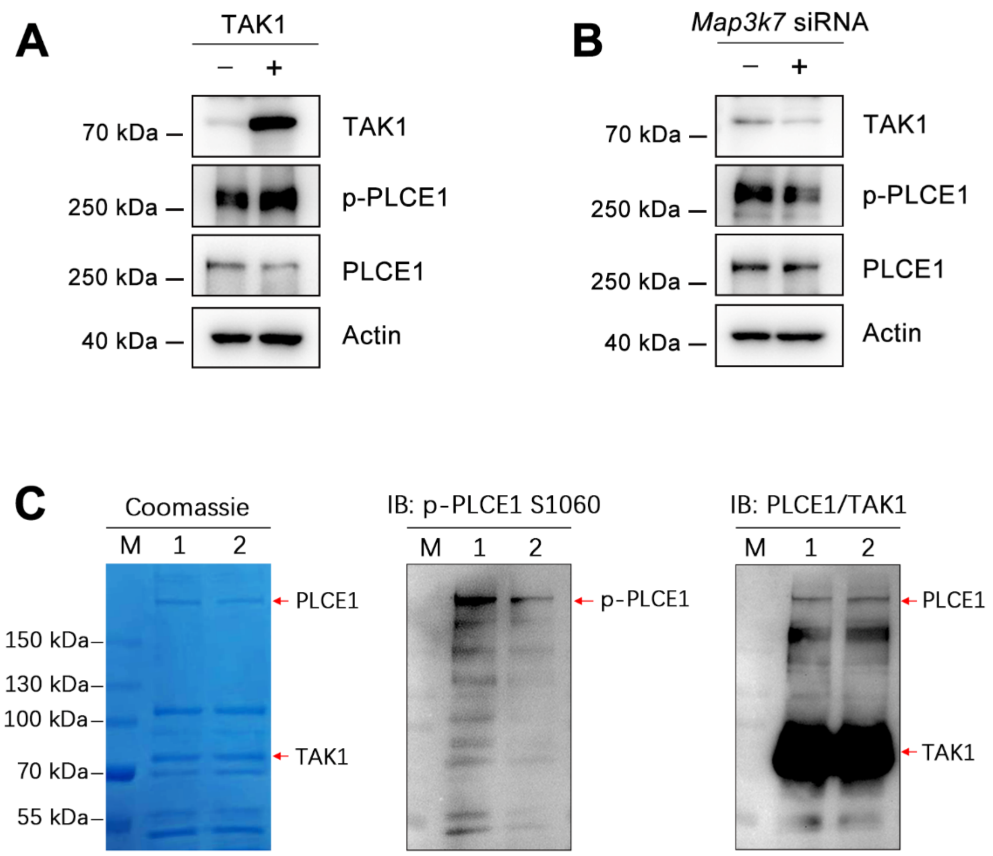


***Figure S6.*** **TAK1 phosphorylates PLCE1 at serine 1060.** A) TAK1 overexpression induces endogenous PLCE1 phosphorylation at serine 1060 (p-PLCE1 S1060). ECA-109 cells were transfected with a plasmid expressing TAK1. 24 h post-transfection, cells were harvested and subjected to western blot analysis. Actin was used as a loading control. B) TAK1 knockdown reduces endogenous p-PLCE1 S1060. ECA-109 cells were transfected with *Map3k7* siRNA. 48 h post-transfection, cells were collected for analyzing p-PLCE1 S1060. Actin was used as a loading control. C) TAK1 phosphorylates PLCE1 at S1060 in an *in vitro* kinase assay. ECA-109 cells were transfected with a plasmid carrying S protein tagged TAK1 (SP-TAK1), Myc tagged PLCE1 (Myc-PLCE1), or Myc tagged PLCE1 S1060A (Myc-PLCE1 S1060A). 24 h post-transfection, cells were harvested and subjected to protein pull-down, the resulting purified SP-TAK1, Myc-PLCE1, and Myc-PLCE1 S1060A were incubated in a kinase assay buffer. The reaction mixtures were then resolved by SDS-PAGE, the resulting gels were analyzed by Coomassie blue staining or western blot. Lane M: marker; Lane 1: SP-TAK1 + Myc-PLCE1; Lane 2: SP-TAK1 + Myc-PLCE1 S1060A. The observed band for p-PLCE1 in lane 2 is likely due to the presence of endogenous wild type PLCE1 in the TAK1 pull-down samples.

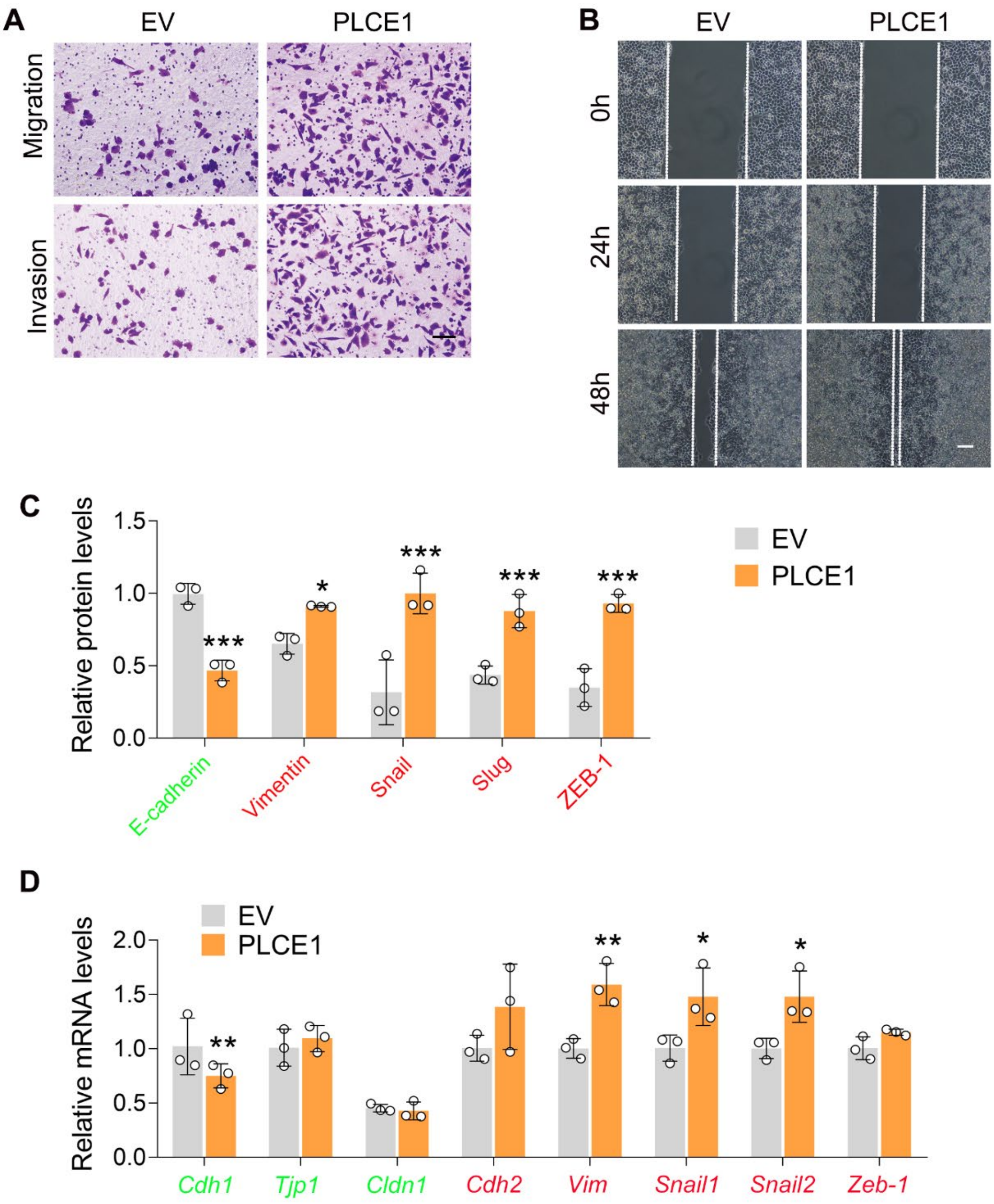


***Figure S7.*** **PLCE1 promotes cell migration and invasion in ECA-109 cells.** A-B) PLCE1 enhances cell migration and invasion in ECA-109 cells (A, transwell assay, Scale bar = 500 μm; B, wound healing assay, Scale bar = 100 μm). C) Q*uantitative analysis of the western blot data as shown in Fig. 3D.* D) PLCE1 increased expression of endogenous mesenchymal markers in ECA-109 cells, while decreased the expression of epithelial markers. Gene expression was analyzed by qRT-PCR. *Gapdh* was used as a house-keeping gene. EV: empty vector. Data are presented as mean ± SD (error bars). Statistical significance was tested by unpaired Student's *t*-test. *n = 3 biologically independent replicates.* *$p < 0.05$, **$p < 0.01$, and ***$p < 0.001$ vs EV.

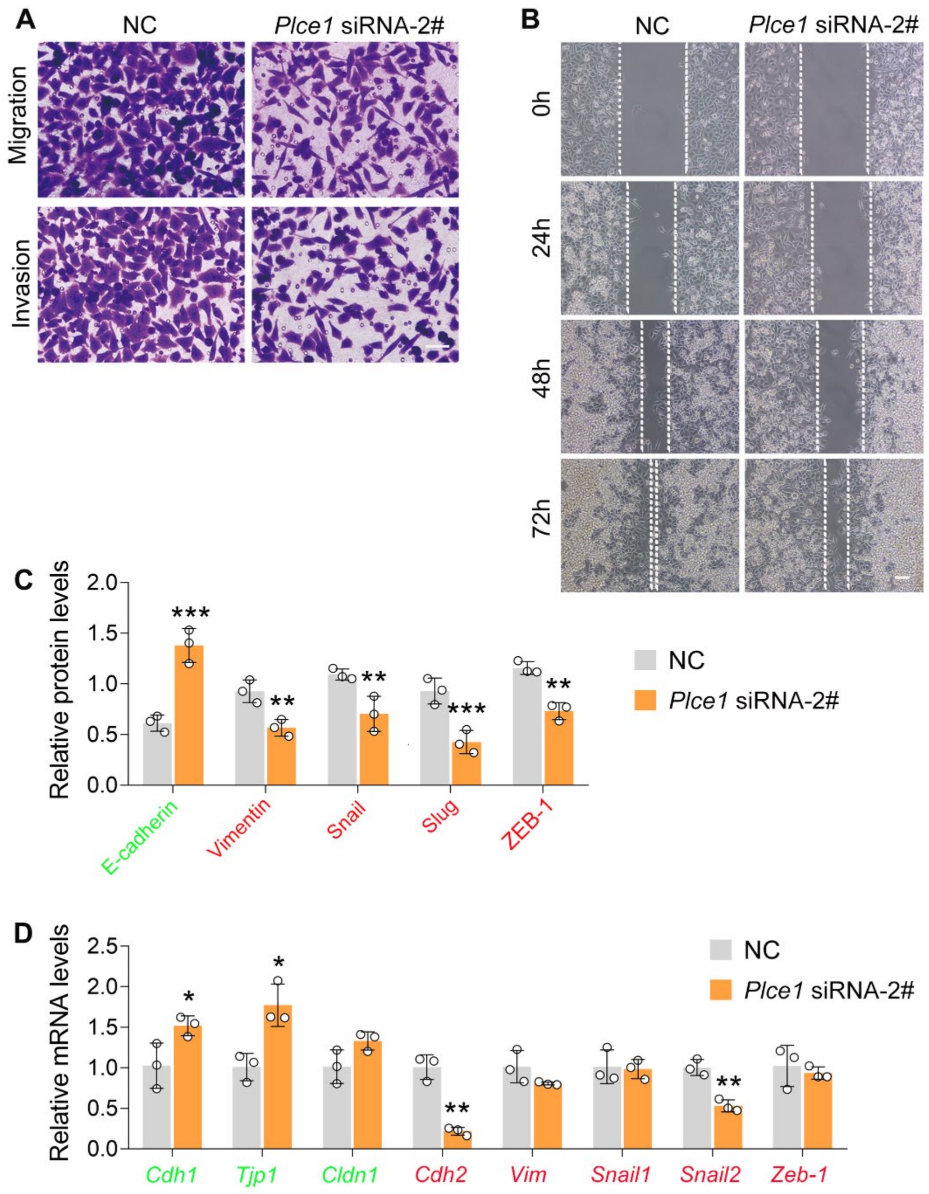


***Figure S8.*** **PLCE1 silencing inhibits cell migration and invasion in ECA-109 cells.** A-B) Reduced expression of PLCE1 inhibits cell migration and invasion in ECA-109 cells. Cell migration and invasion were analyzed by transwell (A) or wound healing assay (B). Scale bar = 50 μm (A) or 100 μm (B). C) The *quantified data of western blots as shown in Fig. 3I.* D) Reduced expression of PLCE1 decreased endogenous mesenchymal marker gene expression in ECA-109 cells, while increased epithelial marker gene expression. Gene expression was assayed by qRT-PCR, and *Gapdh* was used as a house-keeping gene. *NC: negative control.* Data are presented as mean ± SD (error bars). $n$ = 3 biologically independent replicates. Statistical significance was tested by unpaired Student's *t*-test. $^*p < 0.05$, $^{**}p < 0.01$, and $^{***}p < 0.001$ vs NC.

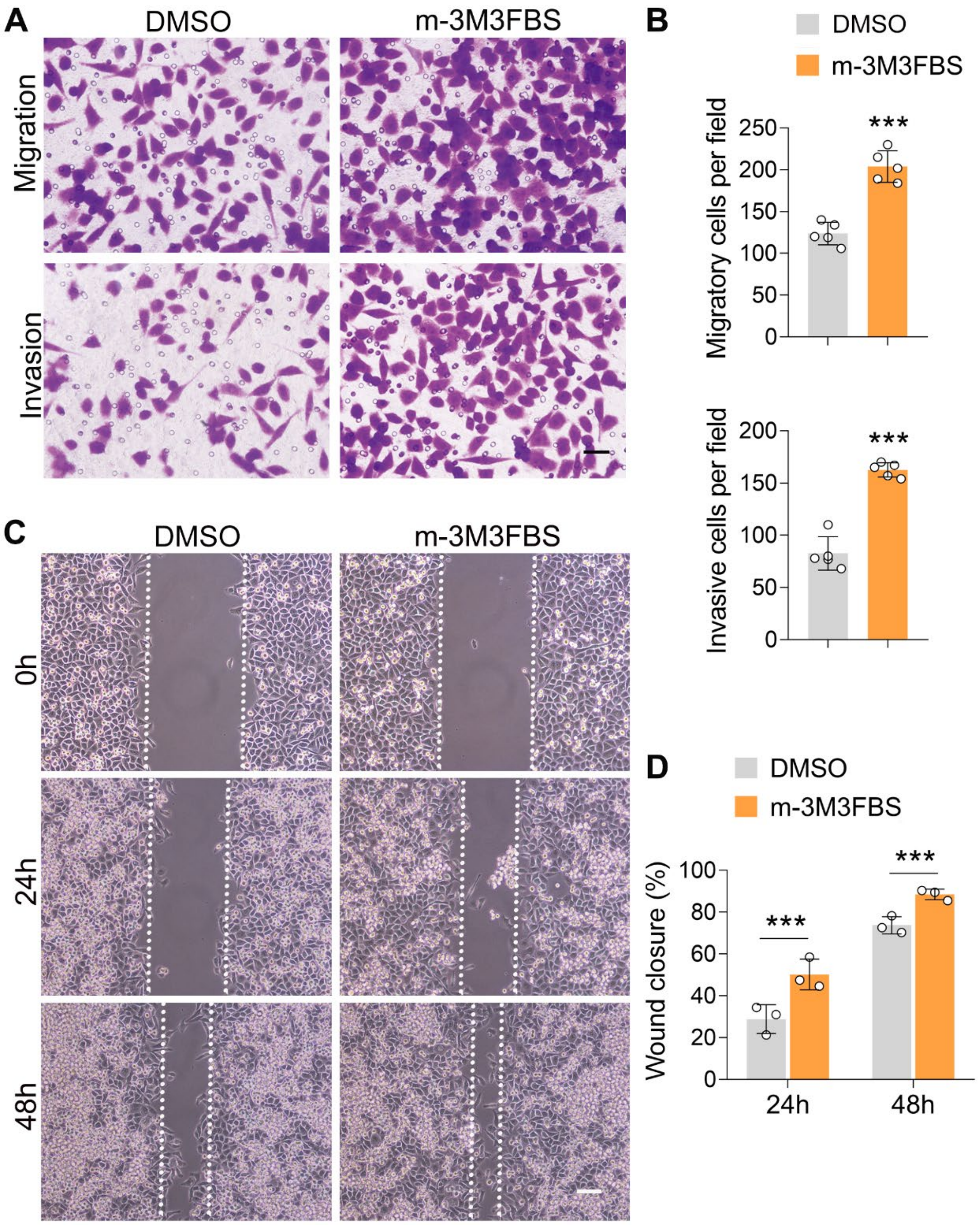


***Figure S9.*** **Activation of PLCE1 stimulates cell migration and invasion in ECA-109 cells.** Cells were treated with 1 μM of m-3M3FBS for 24 h. A-D) Activation of PLCE1 by m-3M3FBS promotes cell migration and invasion (A-B, transwell assay, Scale bar = 50 μm. *n* = 5 biologically independent replicates; C-D, wound healing assay, Scale bar = 100 μm, *n* = 3 biologically independent replicates). Data are presented as mean ± SD. Statistical significance was tested by unpaired Student's *t*-test. ***$p < 0.001$.

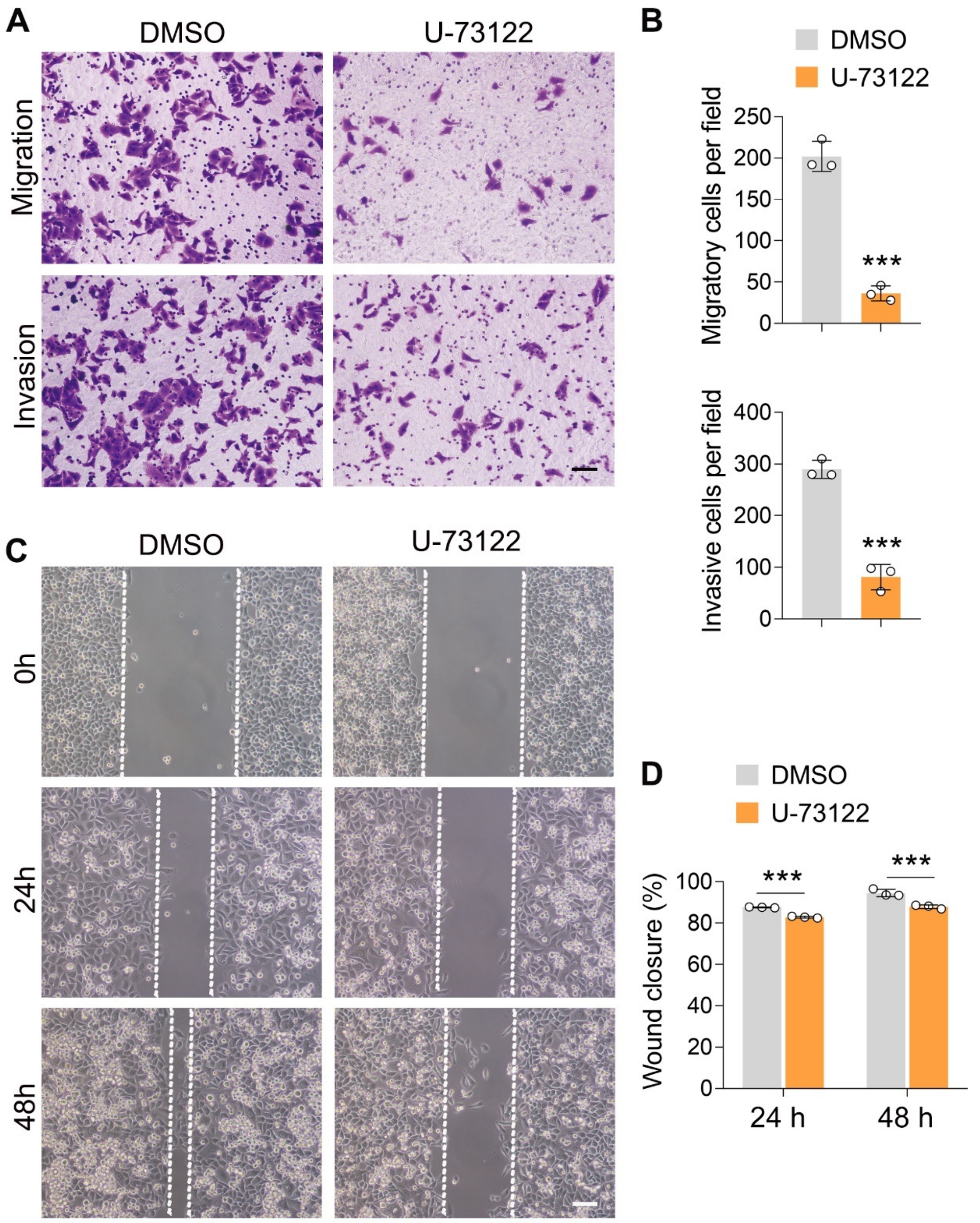


***Figure S10.* Inhibition of PLCE1 reduces cell migration and invasion in ECA-109 cells.** Cells were treated with 10 μM of U-73122 for 24 h. A-D) Inhibition of PLCE1 attenuates cell migration and invasion in ECA-109 cells (A-B, transwell assay, Scale bar = 500 μm; C-D, wound healing assay, Scale bar = 100 μm). $n$ = 3 biologically independent replicates. Data are presented as mean ± SD. Statistical significance was tested by unpaired Student's *t*-test. ***$p < 0.001$ vs DMSO.

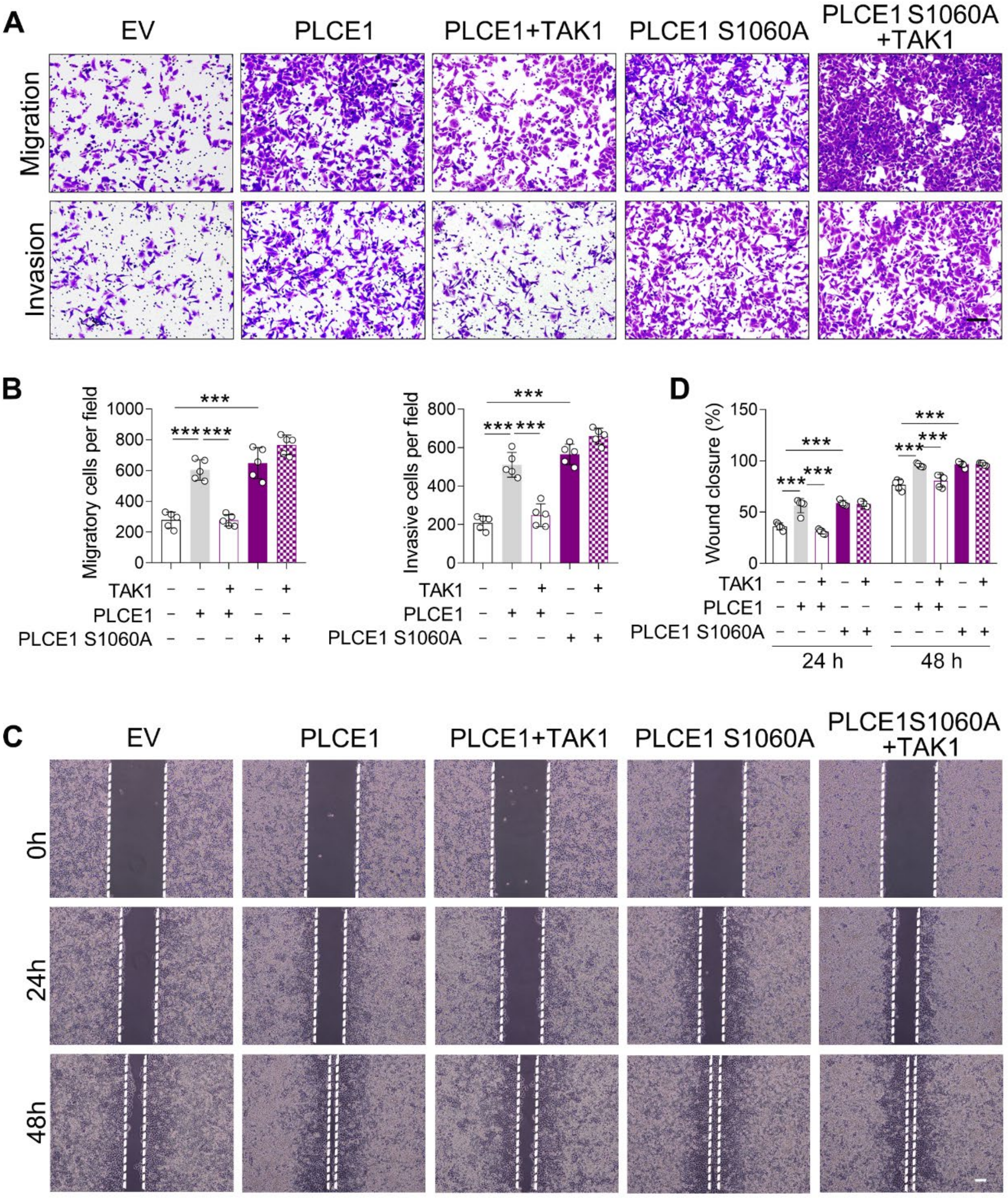


***Figure S11.*** **The negative impact of TAK1 on PLCE1-induced cell migration and invasion requires TAK1 kinase activity.** ECA-109 cells were transfected with plasmids expressing *Plce1*, mutated *Plce1* (S1060A) or *Map3k7* as indicated. 36 h post-transfection, cell migration and invasion were assayed by transwell assay (A-B) and wound healing assay (C-D). $n = 5$ biologically independent replicates. Scale bar = 500 μm (A); Scale bar = 100 μm (C). Data are presented as mean ± SD. Statistical significance was tested by two-tailed one-way ANOVA test. ***$p < 0.001$.

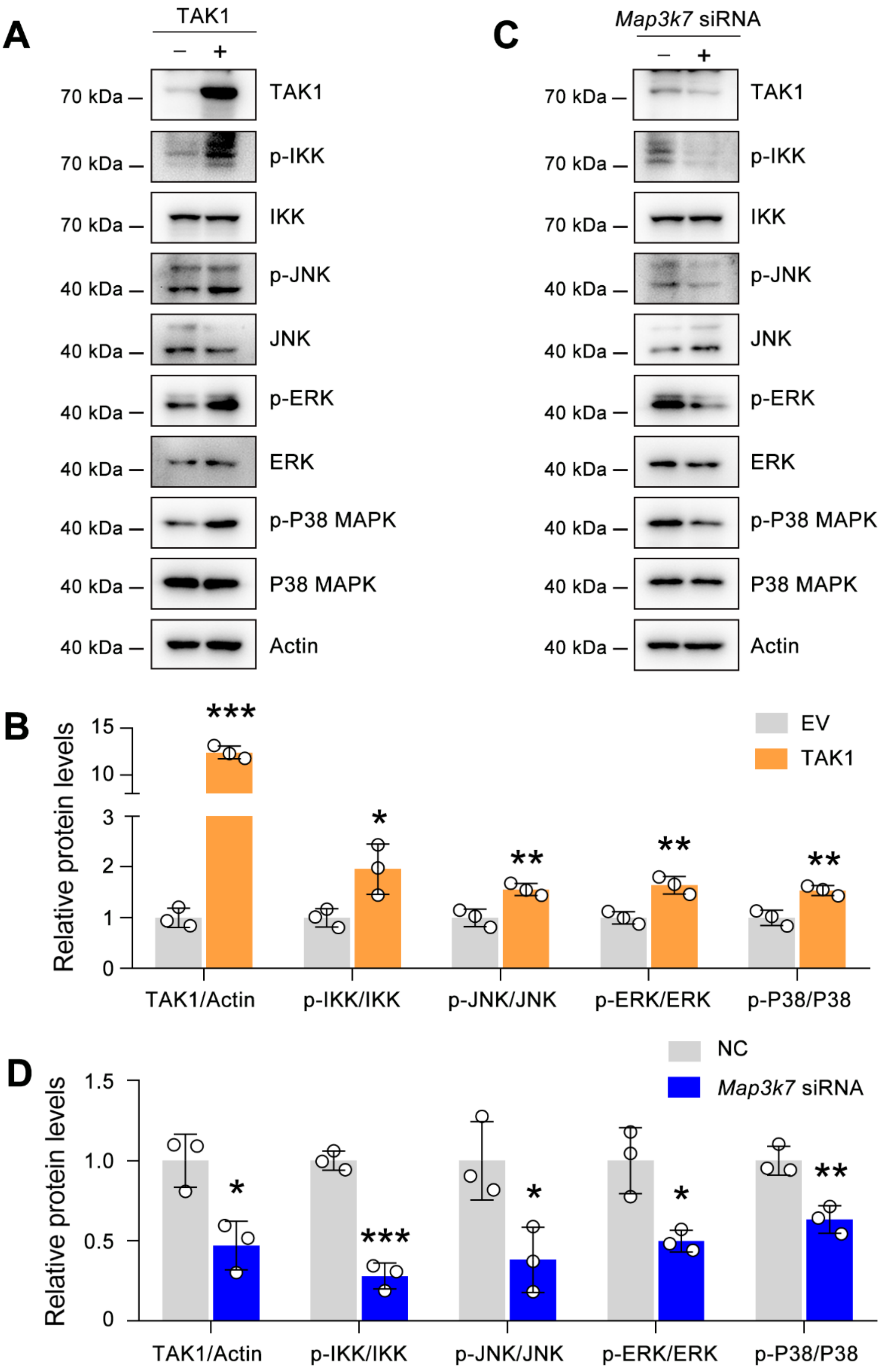


***Figure S12.*** **Effects of TAK1 on p-IKK, p-JNK, p-ERK, and p-P38 MAPK in ECA-109 cells.** A) TAK1 overexpression activates p-IKK, p-JNK, p-ERK, p-P38 MAPK. ECA-109 cells were transfected with a plasmid expressing TAK1. 24 h post-transfection, cells were collected for western blot analysis. B) Quantification data for the blots in (A). C) TAK1 knockdown represses p-IKK, p-JNK, p-ERK, p-P38 MAPK. ECA-109 cells were transfected with *Map3k7* siRNA. 48 h post-transfection, cells were harvested for western blot analysis. D) Quantification data for the blots in (C). EV: empty vector; NC: negative control. Actin was used a loading control. * $p < 0.05$, ** $p < 0.01$, and *** $p < 0.001$, by unpaired Student's *t*-test.

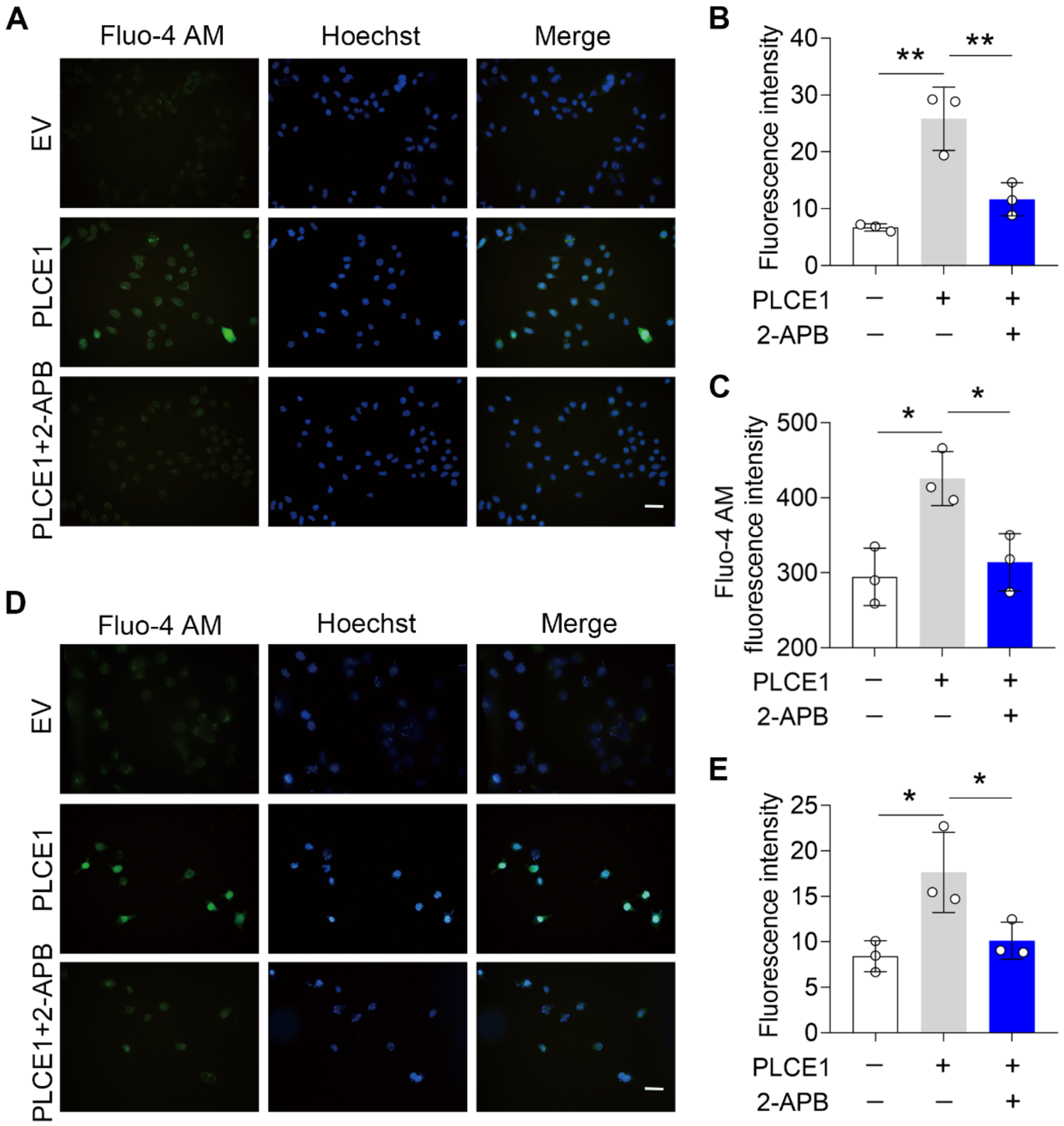


***Figure S13.*** **IP3R blockade inhibits PLCE1-induced intracellular calcium accumulation.** ECA-109 and KYSE-150 cells were transfected with a plasmid expressing *Plce1* for 6 h and then treated with 2-APB (10 μM) for additional 18 h. The intracellular calcium [$Ca^{2+}$] was labeled with Fluo-4 AM. A) Fluorescent imaging the [$Ca^{2+}$] in ECA-109 cells. Scale bar = 20 μm. B) Quantitative analysis of Fluo-4 AM fluorescence intensity as shown in (A). C) Fluorescence intensity of Fluo-4 in ECA-109 cells was examined under an absorbance photometer. $n$ = 3 biologically independent replicates. D) Fluorescent imaging the [$Ca^{2+}$] in KYSE-150 cells. Scale bar = 20 μm. E) Quantitative analysis of Fluo-4 AM fluorescence intensity in KYSE-150 cells as shown in (D). Data are presented as mean ± SD. Statistical significance was tested by two-tailed one-way ANOVA test. $^{*}p < 0.05$ and $^{**}p < 0.01$.

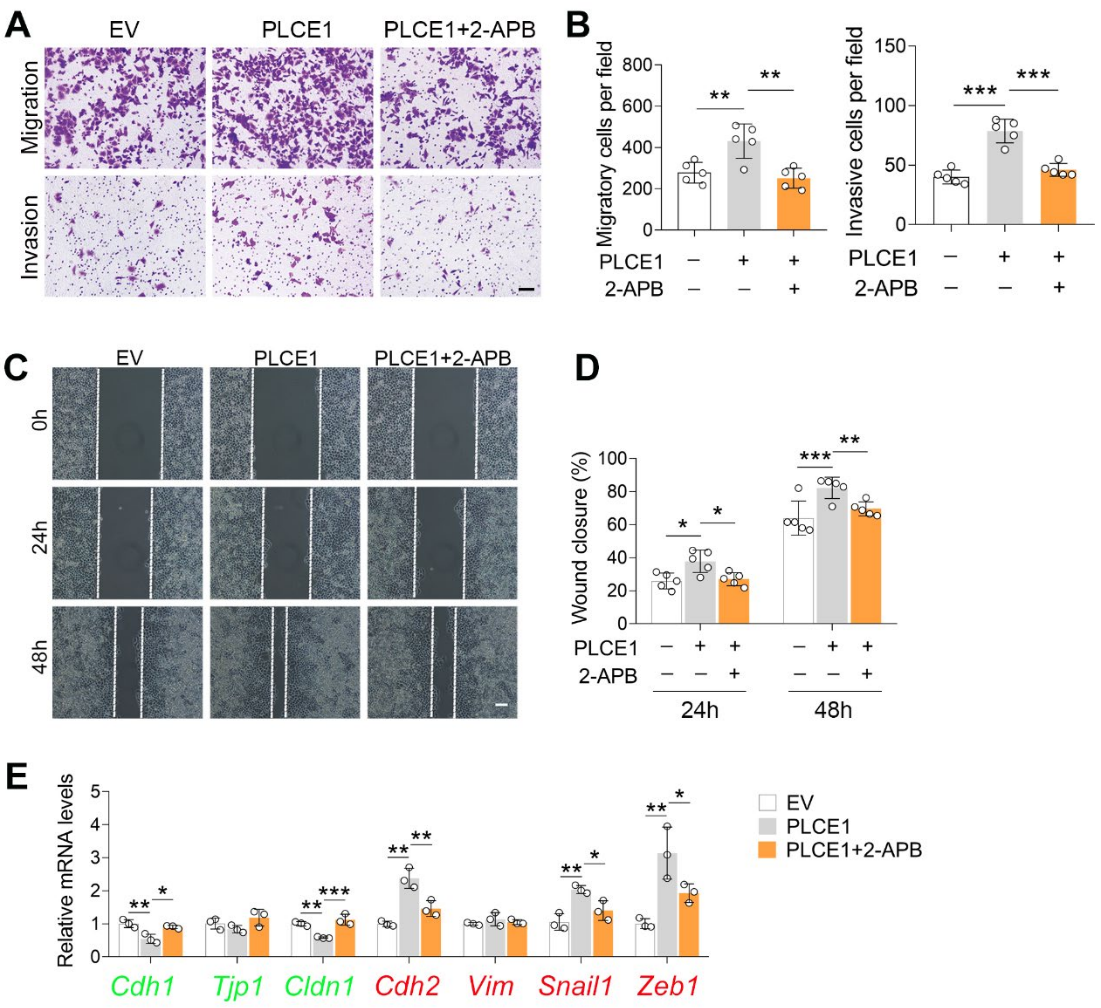


***Figure S14.*** **2-APB treatment counteracts PLCE1-induced cell migration.** A-D) IP3R blocking inhibits PLCE1 promoted cell migration and invasion. ECA-109 cells were transfected with the plasmid expressing *Plce1* for 6 h and then cells were treated with 2-APB (10 μM) for additional 18 h. Cell migration and invasion were analyzed by transwell (A, B) or wound healing (C, D) assay. *n* = 5 biologically independent replicates. Scale bar = 500 μm (A) and 100 μm (C). E, IP3R blocking counteracts PLCE1-induced changes in EMT gene expression. The levels of mRNA were analyzed by qRT-PCR, and *Gapdh* was used as a house-keeping gene. *n* = 3 biologically independent replicates. Data are presented as mean ± SD. Statistical significance was tested by two-tailed one-way ANOVA test. $^{*}p < 0.05$, $^{**}p < 0.01$, and $^{***}p < 0.001$.

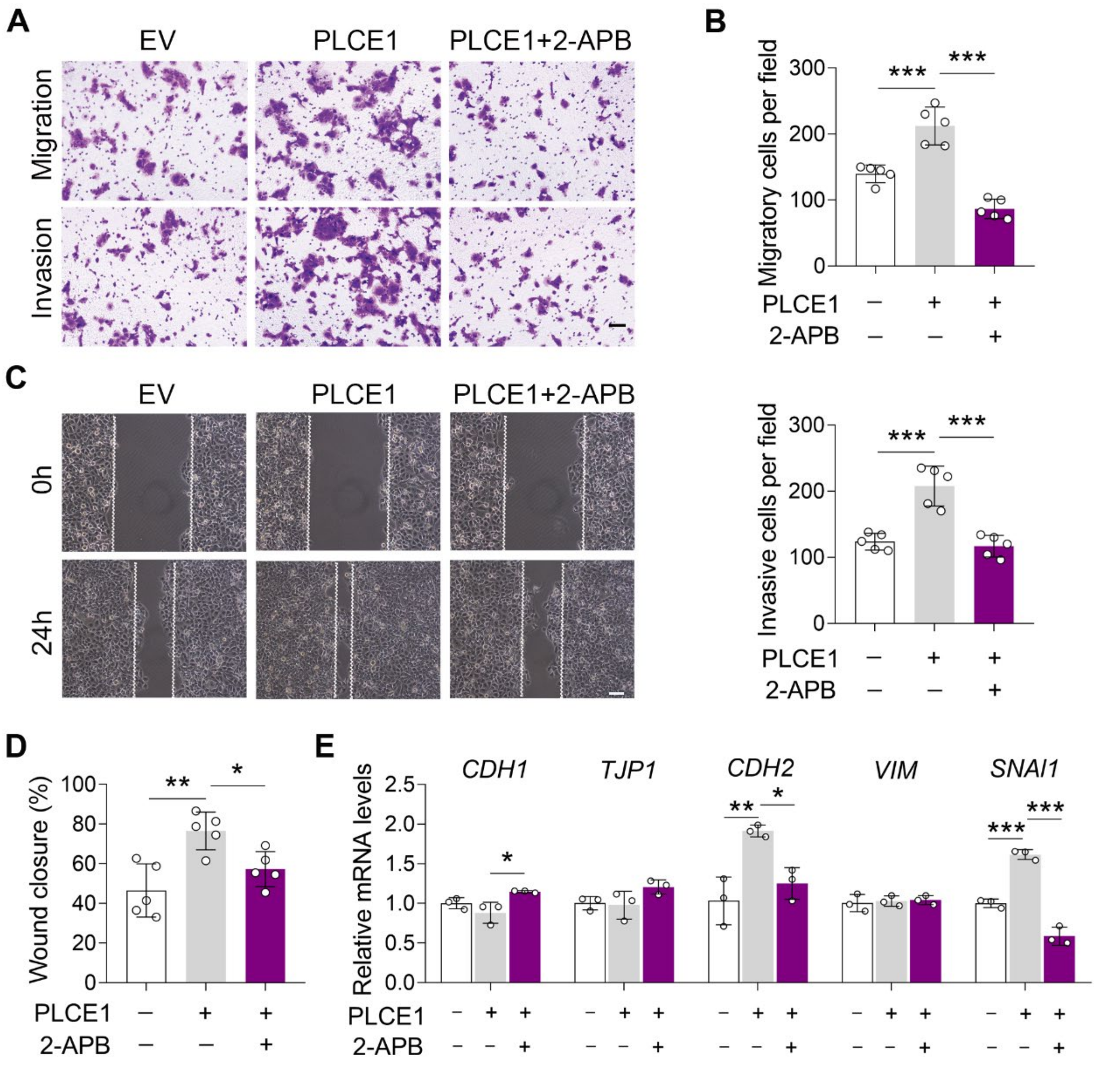


***Figure S15.* IP3R inhibition represses PLCE1-stimulated cell migration and invasion in KYSE-150 cells**. Cells were transfected with plasmid expressing *Plce1* for 6 h and then treated with 2-APB (10 μM) for additional 18 h. A-B) Transwell assay showing the application of 2-APB attenuates cell migration and invasion induced by PLCE1. Scale bar = 500 μm. *n* = 5 biologically independent replicates. C-D) Wound healing assay showing the treatment of 2-APB represses cell migration induced by PLCE1. Scale bar = 100 μm. *n* = 5 biologically independent replicates. E) 2-APB counteracts PLCE1-induced changes in EMT gene expression. The mRNA levels were detected by qRT-PCR, *Gapdh* was used as a house-keeping gene. *n* = 3 biologically independent replicates. Data are presented as mean ± SD. Statistical significance was tested by two-tailed one-way ANOVA test. $^{*}p < 0.05$, $^{**}p < 0.01$, and $^{***}p < 0.001$.

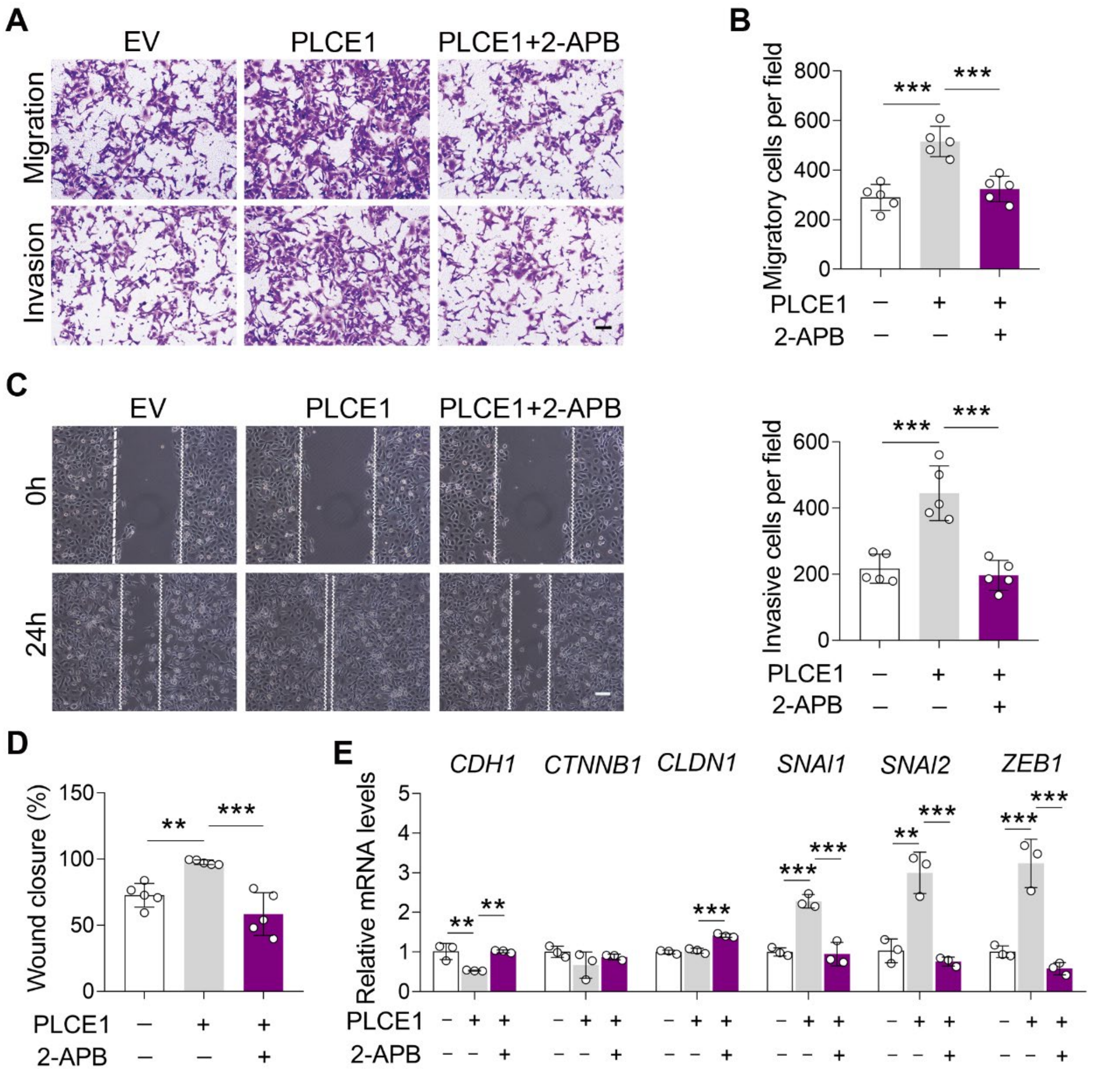


***Figure S16.*** **IP3R inhibition reduces PLCE1-stimulated cell migration and invasion in TE-1 cells**. Cells were transfected with plasmid expressing *Plce1* for 6 h and then treated with 2-APB (10 μM) for additional 18 h. A-D) Cell migration and invasion induced by PLCE1 were repressed by the application of 2-APB in TE-1 cells. Cell migration and invasion were analyzed by transwell assay (A-B, scale bar = 500 μm) and wound healing assay (C-D, scale bar = 100 μm). *n* = 5 biologically independent replicates. E) 2-APB abolishes PLCE1-induced changes in EMT gene expression in TE-1 cells. Gene expression was analyzed by qRT-PCR, and *Gapdh* was used as a house-keeping gene. *n* = 3 biologically independent replicates. Data are presented as mean ± SD. Statistical significance was tested by two-tailed one-way ANOVA test. **$p < 0.01$ and ***$p < 0.001$.

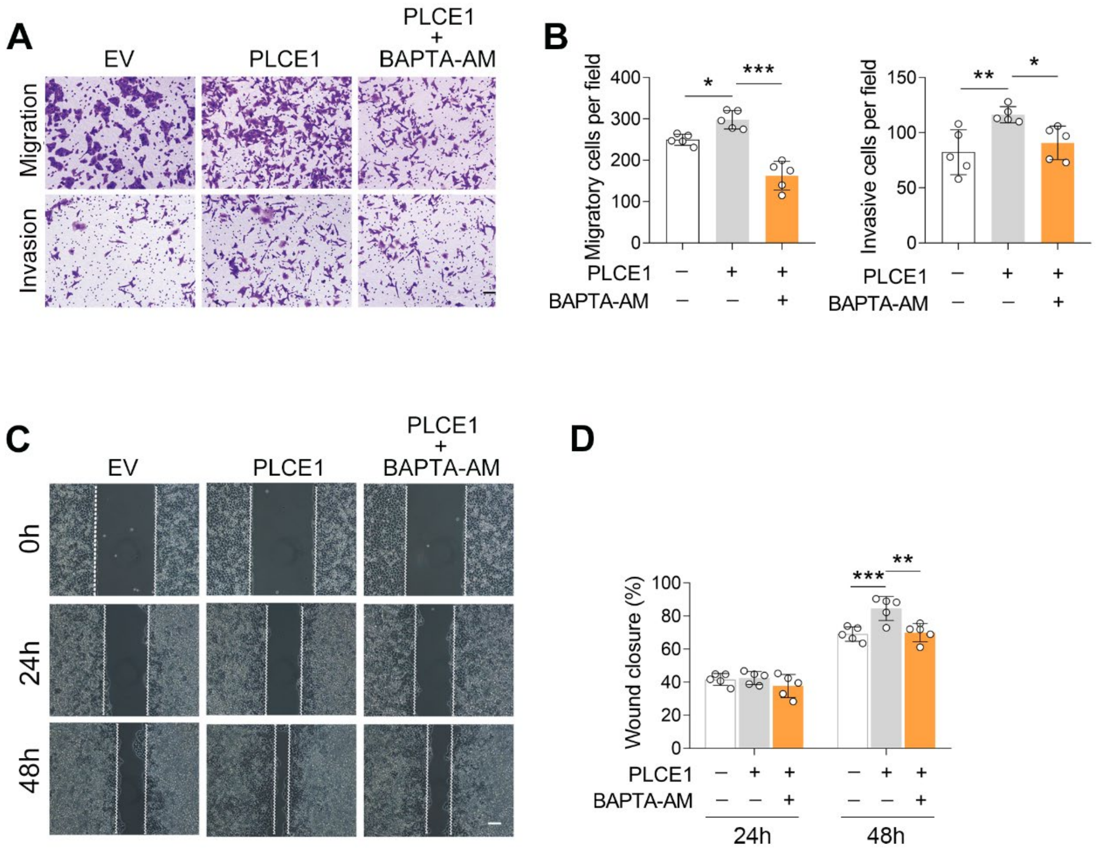


***Figure S17.*** **[$Ca^{2+}$] blockade inhibits PLCE1-induced cell migration and invasion in ECA-109 cells**. A-D) [$Ca^{2+}$] blocking inhibits PLCE1 promoted cell migration and invasion. ECA-109 cells were transfected with the plasmid expressing *Plce1* for 6 h and then treated with BAPTA-AM (10 μM) for additional 18 h. Cell migration and invasion were tested by transwell (A, B) or wound healing (C, D) assay. *n* = 5 biologically independent replicates. Scale bar = 500 μm (A) and 100 μm (C). Data are presented as mean ± SD. Statistical significance was tested by two-tailed one-way ANOVA test. $^{**}p < 0.01$ and $^{***}p < 0.001$.

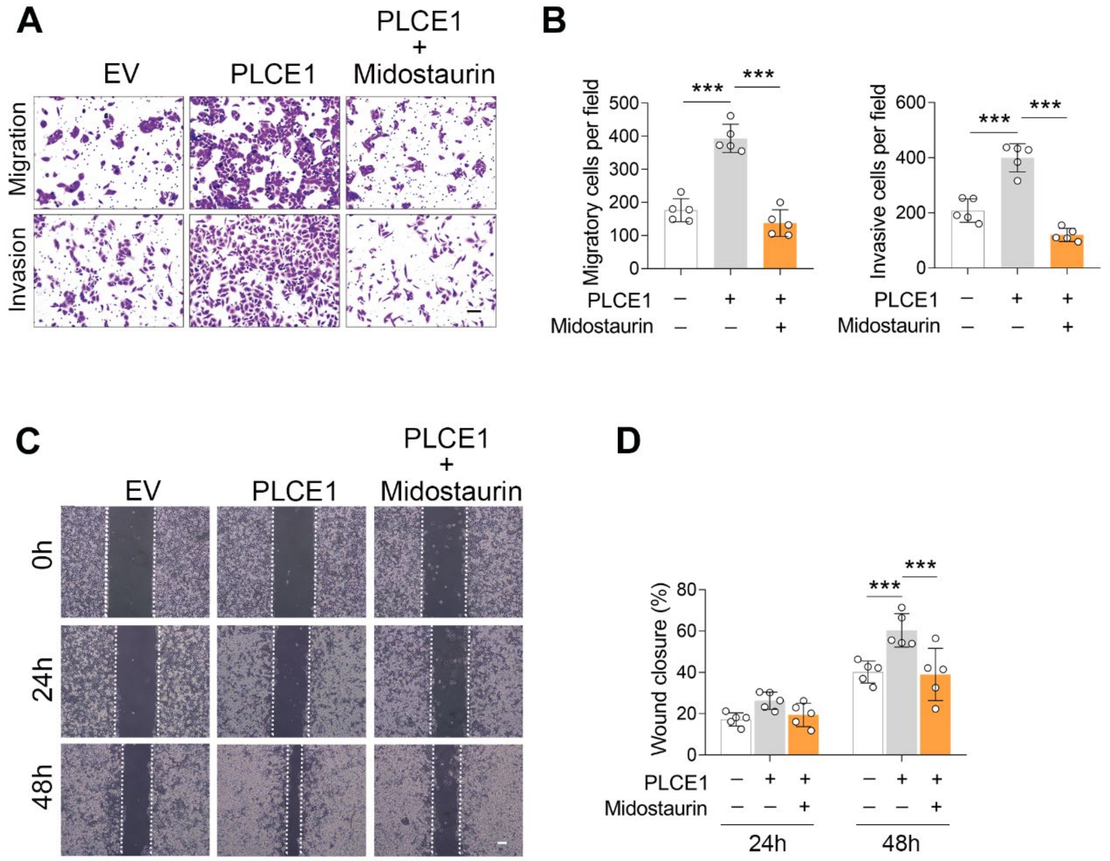


***Figure S18.*** **PKC inhibition counteracts PLCE1-induced cell migration and invasion**. A-D) ECA-109 cells were transfected with the plasmid expressing *Plce1* for 6 h and then treated with Midostaurin (100 nM) for additional 18 h. Cell migration and invasion were tested by transwell (A, B) or wound healing (C, D) assay. *n* = 5 biologically independent replicates. Scale bar = 500 μm (A) and 100 μm (C). Data are presented as mean ± SD. Statistical significance was tested by two-tailed one-way ANOVA test. $^{*}p < 0.05$, $^{**}p < 0.01$, and $^{***}p < 0.001$.

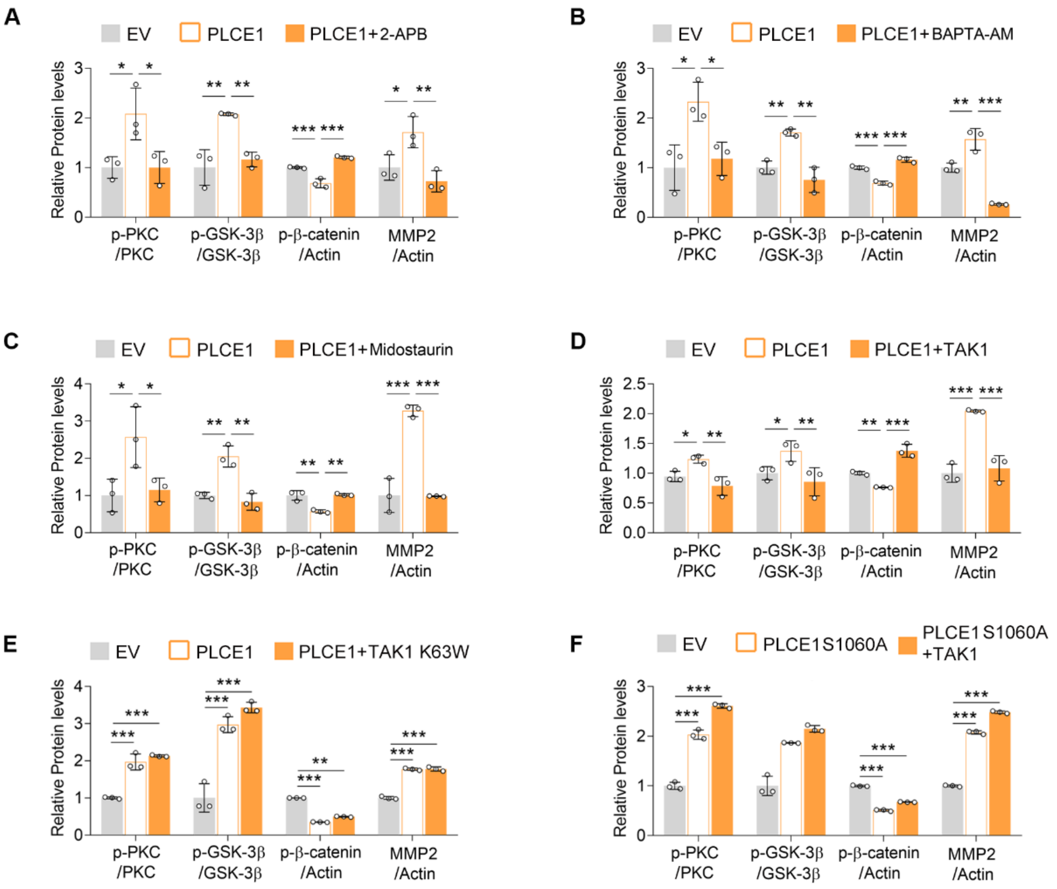


***Figure S19.*** **TAK1 mitigates PLCE1-induced signal transduction in the axis of PKC/GSK-3β/β-Catenin.** A-F) *Quantified data for the western blots as shown in Figure 7. n = 3 biologically independent replicates. Data are presented as mean ± SD.* Statistical significance was tested by two-tailed one-way ANOVA test. **p < 0.05, **p < 0.01,* and ****p < 0.001.*

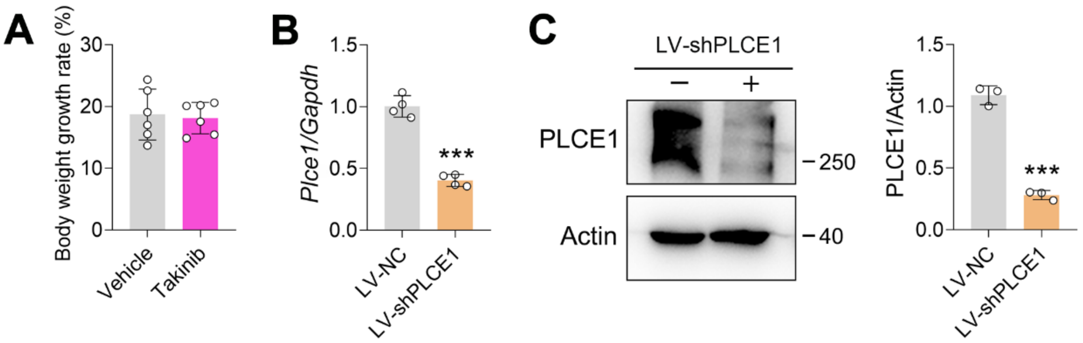


***Figure S20.* The effects of Takinib on mouse body weight and knockdown of PLCE1 by lentivirus in ECA-109 cells.** A) The body weight growth rate was not altered by Takinib. B) The mRNA levels of *Plce1* were decreased by LV-shPLCE1. ECA-109 cells were transduced with LV-shPLCE1 for 48 h, and gene expression was analyzed by qRT-PCR, and *Gapdh* was used as a house-keeping gene. $n$ = 4 biologically independent replicates. C) The protein levels of PLCE1 were decreased by LV-shPLCE1. ECA-109 cells were transduced with LV-shPLCE1 for 48 h, and protein expression was analyzed by western blot, and Actin was used as a loading control. $n$ = 3 biologically independent replicates. ***$p < 0.001$, unpaired Student $t$-test.

**Table S1. The sequences used in gene knockdown and mutation.**

| Genes | Sequences (5'-3') |
|---|---|
| *Map3k7* siRNA | GGAGTTGTTTGCAAAGCTA |
| *Plce1* siRNA-1 | GGACTTCAATATCGCAGTA |
| *Plce1* siRNA-2 | GTCGAAGTGTAGAATTGGA |
| *Plce1* siRNA-3 | CAATCATCATATCGATTGA |
| *Map3k7* gRNA | F: ccgAGGGGCTTCGATCATCTCAC |
| | R: aacGTGAGATGATCGAAGCCCCT |
| *Plce1* (S1060A) | F: TGGAGTGCTCGAAACCCCGCACCCGGAACATCAGCAAA |
| | R: GGGGTTTCGAGCACTCCACCGTCTGCCACCAAACAA |

F: forward; R: reverse.

**Table S2. Primer sequences used in qRT-PCR.**

| *Genes* | Primer sequences (5'-3') |
|---|---|
| *Map3k7* | F: ATTGTAGAGCTTCGGCAGTTATC |
| | R: CTGTAAACACCAACTCATTGCG |
| *Plce1* | F: GGGTGACATGGCTGATCCTC |
| | R: GACAGCGTTGTAGTTGCCCA |
| *Cdh1* | F: GCTTTACTGTTTCTCAAGTGT |
| | R: AATACACAATTATCAGCACCC |
| *Vim* | F: AACTTCTCAGCATCACGAT |
| | R: GTAGGAGTGTCGGTTGTT |
| *Ctnnb1* | F: AGAATTGAGTAATGGTGTAGAAC |
| | R: TACCCATACATATCCCAAATAGT |
| *Cldn1* | F: TGTATAGTCCTCTTGGGTTG |
| | R: AATTGTCAGTGGAGTCAGT |
| *Cdh2* | F: AAAAGGAAAGGAAAGAAAGGG |
| | R: GTCAGAGGTGTATCATTTATATTCT |
| *Zeb1* | F: GTTGCTCCTTCTTCCTGA |
| | R: ATGTGGTTCCTGTTCCTAG |
| *Tjp1* | F: TGTGGACATCCTACTTACTTAA |
| | R: GAGAAGATAAAGAAACTGTTGTATG |
| *Snai1* | F: AGCTATTTCAGOCTCCTG |
| | R: TGTAAACATCTTCCTCOCAG |
| *Snai2* | F: CTGTATGAAACTGAGATGTTGT |
| | R: GAAGCAAGTAAAGTCTCTGAAA |
| *Gapdh* | F: AAGAGCACAAGAGGAAGAG |
| | R: TAACTGGTTGAGCACAGG |

F: forward; R: reverse.

### Materials and methods

#### Antibodies, inhibitors, and growth factors

The following antibodies were used: rabbit polyclonal anti-PLCE1 (#PA5-100856; Invitrogen, Carlsbad, CA, USA). Rabbit monoclonal antibodies against TAK1 (#5206), phospho-TAK1 (Ser412, #9339), PKCα (#2056), phospho-PKC (pan) (gamma Thr514) (#38938), GSK-3β (#9315), phospho-GSK-3β (Ser9) (#5558), β-Catenin (#8480), phospho-β-Catenin (Ser33/37/Thr41) (#9561), MMP-2 (#87809), mouse monoclonal antibodies against Actin (#3700), Myc-Tag (Sepharose® Bead Conjugate) (#55464), Epithelial-Mesenchymal Transition (EMT) antibody sample kit (#9782), phosphor-IKK (#2078), IKK (#61294), phosphor-JNK (#4668), JNK (#9252), phospho-ERK (#4370), ERK (#9102), phospho-P38 MAPK (#9211), P38 MPAK (#9212), and normal rabbit IgG (#2729) were from Cell Signaling Technology (Beverley, MA, USA). TAK1 inhibitors 5Z-7-oxozeaenol (O9890), NG25 (SML1332) and Takinib (SML2216) (#662009) were purchased from Sigma-Aldrich (St. Louis, MO, USA). An intracellular calcium chelator BAPTA-AM (HY-100545), a PKC inhibitor Midostaurin (HY-10230), and an IP3R antagonist (HY-W009724) were obtained from MedChemExpress (New Jersey, USA). EGF Recombinant Human Protein (#PHG0311) was purchased from Gibco (Carlsbad, CA, USA).

#### Plasmid construction

The full-length coding regions of *Map3k7* was synthesized by Heyuan Biotechnology Company (Shanghai, China). The synthesized *Map3k7* was cloned into pcDNA3.1(+) (Invitrogen, Carlsbad, CA, USA) vector using EcoR I and Xho I. Correct construction was confirmed by DNA sequencing. The plasmid of pRK5-N-myc PLCE1 was a gift from Dr. Friedhelm Hildebrandt (1). PLCE1 (S1060A) was generated using a PCR-based mutagenesis kit (Stratagene, La Jolla, CA, USA) using pRK5-N-myc-human PLCE1 as template. The primer sequences are listed in Table S1. All plasmids were confirmed by sequencing. For transient transfection, the plasmids were transfected into ESCC cells with Lipofectamine 2000 (Invitrogen, Carlsbad, CA), according to the manufacturer's instructions.

*Map3k7* short interfering RNAs (siRNAs), *Plce1* siRNAs and a corresponding scrambled siRNA were chemically synthesized at RiboBio (Guangzhou, China). The siRNA sequences are listed in Table S1. The siRNAs were transfected into ECA109 cells with Lipofectamine

RNAiMAX (Invitrogen, Carlsbad, CA), in accordance with the instructions.

**Lentivirus transduction**

LV-*Map3k7* shRNA and LV-NC shRNA were produced at Hanbio (Shanghai, China). LV-*Plce1* shRNA (LV-shPLCE1) and LV-NC shRNA were produced at OBiO (Shanghai, China). Lentivirus transduction was performed as previously described (2). To generate a stale cell line with low PLCE1 expression, ECA-109 cells were transduced with LV-shPLCE1. 24 h post-transduction, cells were treated with 2.5 μg/ml puromycin to eliminate non-transduced cells. The efficiency of lentivirus with knockdown of TAK1 or PLCE1 were verified by qRT-PCR and western blot analysis.

**Quantitative real time-PCR (qRT-PCR)**

Total RNA was isolated from the ESCC cells using TRIzol Reagent (Invitrogen, Carlsbad, CA) and quantified. Then reversely transcribed into First strand cDNA using the PrimeScript RT reagent kit (Takara, Tokyo, Japan) according to the manufacturer's instruction. qRT-PCR was used to analyze the expression level of target gene mRNA by using a Fast Start Master SYBR Green Kit (Roche) on the RocheLightCycler®96 instrument and software (Roche, Basel, Switzerland). The qRT-PCR conditions were as follows: 95°C for 6 min, followed by 40 cycles of 95°C for 10 sec, 60°C for 30 sec, 72°C for 10 sec. Primer sequences were shown in Table S2. Relative quantitation of target genes was normalized to that of GAPDH mRNA, and were analyzed using the $2^{-\triangle\triangle CT}$ method.

**Wound healing assay**

For wound-healing assay, ibidi Culture-Inserts (ibidi GmbH) were used to guarantee the uniformity of each initial (0 h) wound. In brief, $3 \times 10^4$ cells were seeded into the ibidi Culture-Inserts (70 μl cell suspension for each side of the scratch chamber) and incubated in medium containing 10% FBS. After 24 h, cells were grown to almost 100% confluence. Then the scratch chambers were removed to make the wounds and rinsed with PBS for 3 times to remove the suspended cells. Cells were continually cultured in DMEM supplemented with 1%

FBS and photographed randomly at 0 h, 24 h and 48 h, respectively. Wound closure was calculated using the Image J software according to the following formula: Wound healing rate (%) = (24 h or 48 h wound area - 0 h wound area) / 0 h wound area × 100%.

**Transwell assay**

Cell migration and invasion were determined using transwell assay. For migration assay, $5 \times 10^4$ cells in 200 μl of serum-free culture medium were placed into the upper chambers (8.0 μm pore size, Corning, NY) in 24-well plates. For cell invasion assay, $5 \times 10^4$ cells in 200 μl of serum-free culture medium were placed into the upper chambers coated with 80 μl of Matrigel (BD Bioscience). The lower chamber was added with 500 μl of DMEM supplemented with 10% FBS. After incubating for 24 h at 37°C, the non-migrating cells in the upper chamber were gently wiped with a cotton swab. The chambers were then fixed with 4% paraformaldehyde for 20 min, and dyed with 0.1% crystal violet for 30 min. The migration and invasive cells on to the lower side of the chamber were imaged (randomly selected five fields) and counted under a light microscope (Olympus BX51, Tokyo, Japan) at 200X.

**Immunofluorescence**

Cells were seeded into a 24-well plate with glass coverslips ($2 \times 10^4$ per well). Next day, cells were transfected with plasmid expressing PLCE1 or TAK1 for 6 h and then treated with BAPTA-AM (10 μM) or 2-APB (10 μM) or Midostaurin (100 nM) for additional 18 h, respectively. For immunofluorescence staining, cells were fixed with methanol at room temperature for 15-20 min and washed 3 times in PBS, each time for 10 min. Then cells were blocked with 5% BSA solution containing 0.1-0.5% triton X100 for 2 h at room temperature. After blocking, cells were incubated with rabbit anti-β-Catenin antibody overnight at 4°C, followed by an Alexa Fluor 488/594-conjugated secondary antibody incubation for 2 h at room temperature. Finally, the nuclei were stained with Hoechst. Photographs (400X) were taken with an Olympus fluorescence microscope (BX51, Tokyo, Japan).

***In vitro* kinase assay**

Protein pull-down was performed using a previously reported method (2). Briefly, ECA109 cells were transfected with a plasmid expressing Myc tagged PLCE1 (Myc-PLCE1), Myc tagged PLCE1 S1060A (Myc-PLCE1 S1060A), or S protein tagged TAK1 (SP-TAK1). 24 h post-transfection, cells were harvested for preparing total cell lysates, which was then subjected to protein pull-down using the Myc-Tag beads (Sepharose® Bead Conjugate; #55464; CST, Beverley, MA, USA) or S-protein Agarose (#69704; Millipore, Billerica, MA, USA). After 4 h-incubation on a rotator, unbound proteins were removed using ice-cold wash buffer. Myc-PLCE1, Myc-PLCE1 S1060A, and SP-TAK1 were eluted and collected. The purified SP-TAK1 was incubated with Myc-PLCE1 or Myc-PLCE1 S1060A in a kinase assay buffer (Cell Signaling Technology, #9802), in which 200 μM ATP (Cell Signaling Technology, #9804) was added. The reaction was performed at 30°C for 30 min. The reaction was terminated by adding 5 × loading buffer to each sample. After boiling at 100°C for 5 min, the samples were resolved by SDS-PAGE, the resulting gels were analyzed by Coomassie blue stating or western blot.

**SI References**

1. H. Chaib *et al.*, Identification of BRAF as a new interactor of PLCepsilon1, the protein mutated in nephrotic syndrome type 3. *Am J Physiol Renal Physiol* **294**, F93-99 (2008).

2. Shi H*, et al.* (2021) TAK1 Phosphorylates RASSF9 and Inhibits Esophageal Squamous Tumor Cell Proliferation by Targeting the RAS/MEK/ERK Axis. *Adv Sci (Weinh)* 8(5):2001575.
